# Early Lipid Remodeling During Ischemia–Reperfusion Injury Is Associated with Damage- and Protection-Related Lipid Signatures in Acute Kidney Injury

**DOI:** 10.64898/2026.01.26.701694

**Authors:** Fátima Milhano Santos, Paula Cuevas-Delgado, Vanessa Marchant, Lucia Tejedor-Santamaria, Raúl R. Rodrigues-Diez, Adrián M. Ramos, Ana B. Sanz, Alberto Ortiz, Coral Barbas, Marta Ruiz-Ortega

## Abstract

Acute kidney injury (AKI) secondary to kidney ischemia–reperfusion (IRI) is driven by profound metabolic perturbations that shape oxidative stress, inflammation, and cell death responses. Despite growing evidence of lipid dysregulation in AKI, its biological and mechanistic relevance remains unclear. To address this gap, we performed comprehensive untargeted kidney lipidomics using LC-QTOF-MS to delineate dynamic lipid changes during the acute phase of murine kidney IRI. Integration of lipidomic signatures with kidney gene-expression profiling and curated datamining revealed early activation of lipid pathways associated with injury responses. IRI induced marked lipidomic reprogramming, characterized by a marked accumulation of glycerolipids, including triacylglycerols (TGs) species detected exclusively after injury, together with increased levels of sphingolipids (ceramides, sphingomyelins, and hexosylceramides). Cardiolipins and most glycerophospholipids declined sharply following IRI, whereas specific phosphatidylcholines (PCs) exhibited opposite regulation, consistent with dynamic membrane remodeling. Datamining linked TG accumulation and altered transcriptional regulation of PC metabolism to repair-prone type 1 injured proximal tubular cells, highlighting its role in early tubular injury. Correlation analyses revealed strong associations between glycerolipids/sphingolipids and markers of renal dysfunction and inflammation, identifying TG 54:9 as a candidate for injury biomarker. Conversely, PE 40:6, PI 38:6, together with several lysophosphatidylcholines and ether-linked phospholipids, correlated positively with nephroprotective and antioxidant markers. Together, these patterns delineate two major lipid modules: a damage-associated module and a nephroprotection-associated module. These findings establish lipid remodeling as a central yet underexplored determinant of AKI pathogenesis and underscore lipidomics as a powerful discovery tool for identifying novel biomarkers and mechanistic targets in kidney disease.

## Introduction

Acute kidney injury (AKI) is an important clinical challenge associated with high morbidity and mortality worldwide [1,2]. Beyond its immediate impact, AKI significantly increases the risk of chronic kidney disease (CKD), a growing global health burden and major contributor to both all cause and cardiovascular mortality [3]. Among the leading causes of AKI, ischemia–reperfusion injury (IRI) is particularly relevant in kidney transplantation, affecting 30–50% of recipients of deceased donor grafts and frequently resulting in delayed graft function [4,5]. Despite its clinical significance, there are currently no effective therapies for AKI and no reliable biomarkers for early detection, underscoring the urgent need to elucidate the mechanisms underlying this condition.

Lipids are fundamental components of cellular membranes and dynamic regulators of energy metabolism and signaling pathways. Their homeostasis is critical, as dysregulated lipid metabolism is increasingly recognized as a driver of pathological processes [6]. In the kidney, lipid metabolism plays a central role in maintaining energy balance and cellular integrity. Proximal tubular cells, among the most metabolically active cell types, rely predominantly on mitochondrial fatty acid β-oxidation (FAO) to generate ATP and sustain physiological functions [7–9]. During IRI, a transient reduction in blood flow and oxygen availability triggers profound metabolic reprogramming, shifting renal cells toward anaerobic pathways [10,11]. Ischemia impairs enzymatic activity and disrupts cellular metabolism, whereas reperfusion triggers oxidative stress and lipid peroxidation, causing renal damage either directly at the membrane or indirectly through the generation of cytotoxic polyunsaturated lipid-derived aldehydes [10,12]. IRI-induced metabolic adaptations impair FAO and lipid metabolism, reducing the capacity of renal cells to utilize fatty acids for ATP production and ultimately leading to energy depletion. This results in the accumulation of fatty acids and other complex lipids such as triglycerides (TGs), diacylglycerols (DGs), and ceramides, which ultimately drive lipotoxicity and tubular injury and CKD progression [10,11,13]. Understanding the mechanisms underlying lipid dysregulation during IRI is crucial for developing targeted interventions to preserve renal function and prevent progression to CKD. Lipid profiling in IRI offers a unique opportunity to uncover injury pathways driven by metabolic reprogramming and to guide therapeutic strategies aimed at mitigating cell death and organ damage.

Over the past five decades, most studies have relied on traditional lipid panels, measuring total cholesterol, lipoproteins, and TGs, without capturing the full chemical diversity of lipids [14,15]. Recent advances in analytical mass spectrometry have transformed this landscape, enabling comprehensive profiling of the tissue and biological fluid lipidome. Lipidomic approaches provide an unbiased, high-resolution analysis of small molecules, offering a powerful platform for their identification and quantification in biological systems. By delivering a hypothesis-free metabolic “snapshot” of tissues, these techniques reveal physiological and pathological states, uncovering unexpected metabolic shifts that yield critical insights into complex diseases, including kidney disorders [15–17]. Despite these advances, most clinical and preclinical lipidomics studies have focused predominantly on circulating lipids, offering only a limited view of kidney-specific alterations. Moreover, integration with other molecular layers and correlation with disease-relevant parameters remain insufficiently explored [15]. In this study, we aimed to comprehensively characterize kidney lipidomic alterations during the acute phase of IRI-induced AKI and to evaluate their associations with established markers of kidney damage, functional parameters, and nephroprotective factors. To this end, we applied an untargeted lipidomic approach using reversed-phase liquid chromatography coupled to quadrupole time-of-flight mass spectrometry (LC-QTOF-MS), integrating lipidomic signatures with renal gene expression profiling and curated data mining. This strategy provides novel insights into the lipidomic remodeling that accompanies AKI and highlights its potential mechanistic and translational relevance.

## Materials and Methods

### Kidney Ischemia–Reperfusion Injury Model

Animal procedures were performed following European Community and ARRIVE guidelines for animal experiments, with the prior consent of the Experimental Animal Ethics Committee of the Health Research Institute of the IIS-Fundación Jiménez Díaz and PROEX 065/18 and PROEX 242.2/21 of the Comunidad de Madrid. Bilateral kidney IRI was induced using a dorsal surgical approach, as previously described [18]. Both renal pedicles were clamped for 45 minutes using atraumatic microvascular clamps. Successful induction of ischemia and reperfusion was confirmed by kidney color change, and animals without these visual changes were excluded. Sham-operated mice underwent the same procedure, but without pedicle clamping. All procedures were performed under 2-3% isoflurane anesthesia, maintaining body temperature at 37°C. Mice were euthanized 24h post-surgery, and the kidneys and blood were collected for further analysis. Kidneys were immediately frozen in liquid nitrogen and stored until further analysis. Serum was stored at −80 °C until further biochemical analysis. Serum BUN (blood urea nitrogen) and creatinine levels were measured at IIS-FJD using a Cobas C702 module (Roche Diagnostics, Rotkreuz, Switzerland) according to the manufacturer’s instructions.

### Lipidomics analysis

#### Reagents

High-purity solvents suitable for LC-MS analyses, including methanol (MeOH), acetonitrile (ACN), chloroform (CHCl_3_) and isopropanol (IPA), were obtained from Fisher Scientific (Pennsylvania, USA). Ammonium fluoride (NH4F) (ACS reagent, ≥98%) was purchased from Sigma-Aldrich (Steinheim, Germany). Ammonia solution (28%, GPR RECTAPUR®) and glacial acetic acid (AnalaR® NORMAPUR®) were supplied by VWR Chemicals (Pennsylvania, USA). Formic acid 98% of mass spectrometry grade was from Honeywell International Inc. (Muskegon, USA). Reversed-osmosed ultrapure water was produced using a Milli-Q system (Millipore, Billerica, MA, USA) and was utilized in all aqueous preparations. SPLASH™ LIPIDOMIX™ Quantitative Mass Spec primary standard was obtained from Avanti Polar Lipids (USA).

#### Sample preparation

Tissue disruption and homogenization were performed as previously described [19], with modifications to adapt the procedure for compound extraction using the Folch method [20]. A complete description of the methodology is provided in the Supplementary Information. Briefly, kidney tissue (∼20 mg) was homogenized with methanol (MeOH) with two steel beads (2.8 mm mean diameter) using a TissueLyser LT bead-mill homogenizer (QIAGEN, Hilden, Germany). The homogenate was transferred to glass tubes and subjected to chloroform-based extraction using a CHCl_3_/MeOH/H_2_O ratio of 8:4:3 (v/v). After centrifugation, the lower organic phase was collected for lipidomics analysis and spiked with Splash Lipidomix™ (Sigma-Aldrich) as an internal standard. Quality control (QC) samples were generated by combining equal aliquots from all individual samples. Before sample extraction and instrumental analysis, samples were randomized to minimize potential batch effects.

#### Untargeted lipidomics analysis by UHPLC-ESI-QTOF-MS

The LC-MS analysis of the kidney extracts was conducted as previously described [21–23], using an Agilent 1290 Infinity II UHPLC system coupled to a 6546-quadrupole time-of-flight (QTOF) mass spectrometer (Agilent Technologies) in both positive and negative ion modes to ensure the detection of a broad range of lipid compounds. Samples (1 μL) were injected using an Agilent 1290 Infinity II Multisampler system with a multi-wash option. Reversed-phase chromatography was performed using an InfinityLab Poroshell 120 EC-C8 (3.0 × 100 mm, 2.7 µm) (Agilent Technologies) column, equipped with an appropriate guard column (Agilent InfinityLab Poroshell 120 EC-C18, 3.0 × 5 mm, µm), maintained at 50 °C. The mobile phases for both ionization modes consisted of solvent A) 10 mM ammonium acetate, 0.2 mM ammonium fluoride in 9:1 water/methanol, and solvent B) 10 mM ammonium acetate, 0.2 mM ammonium fluoride in 2:3:5 acetonitrile/methanol/isopropanol mixture. Mass spectrometry analysis was performed using a dual AJS ESI ion source under the following parameters: 175 V fragmentor, 65 V skimmer, 3500 V capillary voltage, 750 V octopole radio frequency voltage, 10 L/min nebulizer gas flow, 200°C gas temperature, 40 psi nebulizer gas pressure, 12 L/min sheath gas flow, and 300°C sheath gas temperature. Data were acquired in positive and negative ESI modes using full scan mode (m/z range 100-1700 at a scan rate of 3 spectra/s). The MS and MS/MS scan rates were set to 3 spectra/s (m/z range 40–1700) with a narrow MS/MS isolation width (∼1.3 amu), and a selection of three precursors per cycle. The MS/MS threshold was set at 5000 counts and 0.001%. Five iterative-MS/MS runs were performed with a collision energy of 20 eV, followed by five additional runs at 40 eV.

#### Data processing and analysis

Raw data obtained from the untargeted UHPLC–MS lipidomics analysis were pre-processed to eliminate background signals and unrelated ions using the recursive workflow in MassHunter Profinder software (version B.10.0.2, Agilent Technologies, Santa Clara, CA, USA). The first step involved chromatographic deconvolution using the Molecular Feature Extraction (MFE) algorithm, which grouped related ions into molecular features based on coelution patterns, charge states, isotopic distributions, and the presence of adducts or dimers. In parallel, features were aligned across all samples based on accurate mass and retention time (RT), generating a consensus spectrum per compound. These aligned features were then subjected to recursive extraction using the Find-by-Ion (FbI) algorithm, which refined the feature list by applying median mass, RT, and consensus spectra to enhance detection consistency. To improve adduct annotation, the following ions were considered: [M + H]+, [M + Na]+, [M + K]+, [M + NH_4_]+ and [M + C_2_H_6_N_2_ + H]+ in positive mode; [M − H]−, [M + Cl]−, [M + CH_3_COOH − H]−, and [M + CH_3_COONa − H]− in negative mode. Neutral loss of water (H_2_O) was also accounted for in both ionization modes [21,22]. The quality of chromatograms and feature consistency were evaluated using MassHunter Qualitative Analysis (version B.10.00, Agilent Technologies). Additionally, Hotelling’s T^2^ range plots from principal component analysis (PCA-X) models were used to assess sample variability, and control charts were constructed by plotting total ion signal intensities versus acquisition order to monitor analytical precision [17,19]. Prior to statistical analysis, the dataset underwent normalization and feature selection. First, the coefficient of variation (CV) of internal standards (IS) was calculated, and features were filtered based on their reproducibility in QC samples, applying a CV threshold of 20% for LC–MS data. Features with signal intensities in blanks exceeding 10% of their average signal in samples were excluded. Missing values were imputed using the K-nearest neighbors (kNN) algorithm. Given the heterogeneity and fibrotic content in renal tissue, a group-wise quantile normalization strategy was implemented to minimize unwanted intra-group biological variability [19]. After preprocessing, both univariate (UVA) and multivariate (MVA) analyses were conducted to explore group differences. For MVA, the data matrix was imported into SIMCA P+16 (Umetrics®, Umea, Sweden), and unsupervised PCA-X as well as supervised partial least squares discriminant analysis (PLS-DA) models were constructed. Metabolites with a variable importance in projection (VIP) ≥ 1, an absolute correlation coefficient |p(corr)| ≥ 0.5, and jackknife confidence intervals excluding zero were considered significant. All PLS-DA models were cross-validated and assessed using CV-ANOVA tool to ensure model robustness [17]. For UVA, MATLAB R2018a (MathWorks, Natick, MA, USA) was used. Group comparisons (cases vs controls) were assessed with the Mann–Whitney U test, and multiple testing correction was applied using the Benjamini–Hochberg method to control the false discovery rate (FDR) at α = 0.05. For each feature, percent change (%Change), fold change (FC), and log2 fold change (Log_2_FC) were calculated to evaluate the magnitude and directionality of differences [17]. Lipid annotation followed a three-step workflow: (i) tentative identification based on MS1 data using the CEU Mass Mediator online tool (http://ceumass.eps.uspceu.es/mediator/), (ii) reprocessing of raw LC–MS/MS data with Lipid Annotator software (Agilent Technologies Inc.), and (iii) manual validation of MS/MS spectra using MassHunter Qualitative Analysis software (version 10.0) by comparing RTs and fragmentation patterns with spectral libraries [21–23]. Lipid nomenclature followed the latest shorthand annotation recommendations [24]. Once the most relevant lipids were identified, heatmaps with hierarchical clustering were generated in MetaboAnalyst 6.0 (https://www.metaboanalyst.ca/) [25], using Euclidean distance and Ward’s clustering method.

#### Statistical analysis

A supervised multivariate model was built to highlight sample clustering and explore lipid associations with experimental conditions. The values of R^2^ = 0.646 and Q^2^ = 0.565 obtained from the PLS-DA model indicated a good to moderate goodness-of-fit and good predictive ability. PLS-DA models for pairwise group comparisons were created to identify lipid features that contributed most to the discrimination between experimental groups. Based on the PLS-DA models, VIP, JK and *p*(corr) values were calculated to evaluate the statistical significance of the compounds. PLS-DA score plots indicate that the kidney lipidomic profile changes significantly following IRI (**Supplementary Figure 1A**). Based on UVA and FDR correction for multiple hypothesis testing (Benjamini–Hochberg method), compounds showing significantly altered abundance (adjusted p-value ≤0.05) in IRI compared to undamaged controls were listed (**Supplementary Table 1**).

### Gene expression analysis by Quantitative Real-Time PCR

#### RNA extraction and analysis

Full RNA was extracted from kidney samples using TRItidy GTM (PanReac, Barcelona, Spain) and quantified using a nanodrop. cDNA was synthesized from 2 μg of total RNA using a High-Capacity cDNA Archive kit (Applied Biosystems) following the manufacturer’s instructions. Quantitative gene expression analysis was performed on a QuantStudio™ 3 Real-Time PCR System (Applied Biosystems) using fluorogenic TaqMan assays (Thermo Fisher Scientific) or predesigned qPCR assays (Integrated DNA Technologies, IDT) with the Premix Ex Taq PCR master mix (Takara), according to the manufacturer’s instructions. Assays used are provided in Table 1 as supplementary information.

#### Data analysis

Relative quantification analysis was performed using QuantStudio design and analysis software based on mRNA copy numbers (Ct value). Results were expressed as n-fold compared to the control group considering the mean ± standard error of the mean (SEM) of 6 animals per group. Data were normalized considering the expression values of Ppia (Mm.PT.39a.2.gs). Statistical analyses were performed using GraphPad Prism version 9.4.1 (GraphPad Software, San Diego, CA, USA). Data distribution and variance were first assessed to guide the choice of statistical tests. Student’s t-test or Welch’s t-test was applied for normally distributed data, depending on variance equality, whereas the Mann–Whitney U test was used for non-normally distributed data. Heatmaps and correlation analysis were performed using MetaboAnalyst 6.0 [25]. Data were log_2_-transformed for normalization, and correlations were assessed using Pearson correlation.

#### RNA-Seq dataset and lipid-focused functional analysis

Publicly available RNA-Seq data deposited in the NCBI Gene Expression Omnibus (GEO) database (accession number GSE186316) were retrieved from our previous paper [26]. We next evaluated early transcriptomic changes in mouse kidneys 4 hours after IRI to identify molecular events that may precede the lipid alterations observed at 24h in this study. Differentially expressed genes were extracted by comparing kidneys 4 h post-IRI (IRI_4h) with controls without injury (Ctrl_4h). Genes with a false discovery rate (FDR) < 0.05 were considered significant and retained for downstream functional analyses. To explore biological pathways specifically related to lipid metabolism, functional enrichment analysis was performed using STRING (Search Tool for the Retrieval of Interacting Genes/Proteins; https://string-db.org/, version 12.0) [27]. For biological interpretation, enriched and Reactome terms were filtered to retain pathways associated with lipid metabolic process (GO:0006629) and Cellular lipid metabolic process (GO:0044255).

#### Data mining of kidney single-cell transcriptomic datasets

To contextualize the predicted lipidomic signatures at the cellular level, single-cell transcriptomic data were explored using the KIT (Kidney Interactive Transcriptomics) portal, a public interactive resource provided by the Humphreys Laboratory (https://humphreyslab.com/SingleCell/). This platform aggregates multiple high-resolution single-cell atlas datasets generated from the mouse or human kidney. Analyses were conducted using the reference dataset “Mouse UUO + IRI sci-RNA-seq3” [28], which provides high-resolution single-cell transcriptional profiles across multiple kidney cell populations under obstructive and ischemic injury. Genes of interest were queried using the “Search for clustering of genes” function, and expression patterns were visualized across annotated kidney cell types. These expression distribution plots were used qualitatively to determine the predominant cellular compartments expressing lipid-related genes, providing cellular localization and contextual interpretation.

## Results

### Ischemia–reperfusion injury induces profound changes in the kidney lipidomic profile

The kidney lipidomic profile assessed by untargeted lipidomics using LC-QTOF-MS disclosed that 80 compounds showed significantly altered abundance (adjusted p-value ≤0.05) in kidney IRI compared to undamaged control kidneys (**Supplementary Table 1**). IRI induced changes in the levels of several lipid classes, including triglycerides (triacylglycerols, TG), diglycerides (diacylglycerols, DG), monoacyl/diacyl-glycerophosphocholines (LPC/PC), diacyl-glycerophosphoserines (PS), and diacylglycerophosphoinositols (PI), among others (**Figure 1A and 1B**). Among these lipids, TGs were the most significantly altered lipids, with levels markedly increased in response to IRI. Indeed, some TG species were detected exclusively after ischemic injury. DG and several classes of sphingolipids, including ceramides, sphingomyelins (SM), and hexosylceramides (HexCer), were also significantly increased. Conversely, most fatty acyl carnitines (CAR) and glycerophospholipids, including some phosphatidylcholines (glycerophosphocholines, PC), were significantly reduced following IRI, with a consistent decrease in the levels of all identified cardiolipins (CL), lysophosphatidylcholines (monoacylglycerophosphocholines, LPC), and ether-linked glycerophosphocholines (Ether-linked glycerophosphocholines, EtherPC). Nevertheless, some specific species of PCs, CAR, phosphatidylethanolamines (diacylglycerophosphoethanolamines, PE), and Ether-linked glycerophosphoethanolamines (EtherPE) exhibited significant increases after kidney injury, including CAR 16:0;O, PE(34:3), and EtherPE(38:6e).

**Figure 1.**
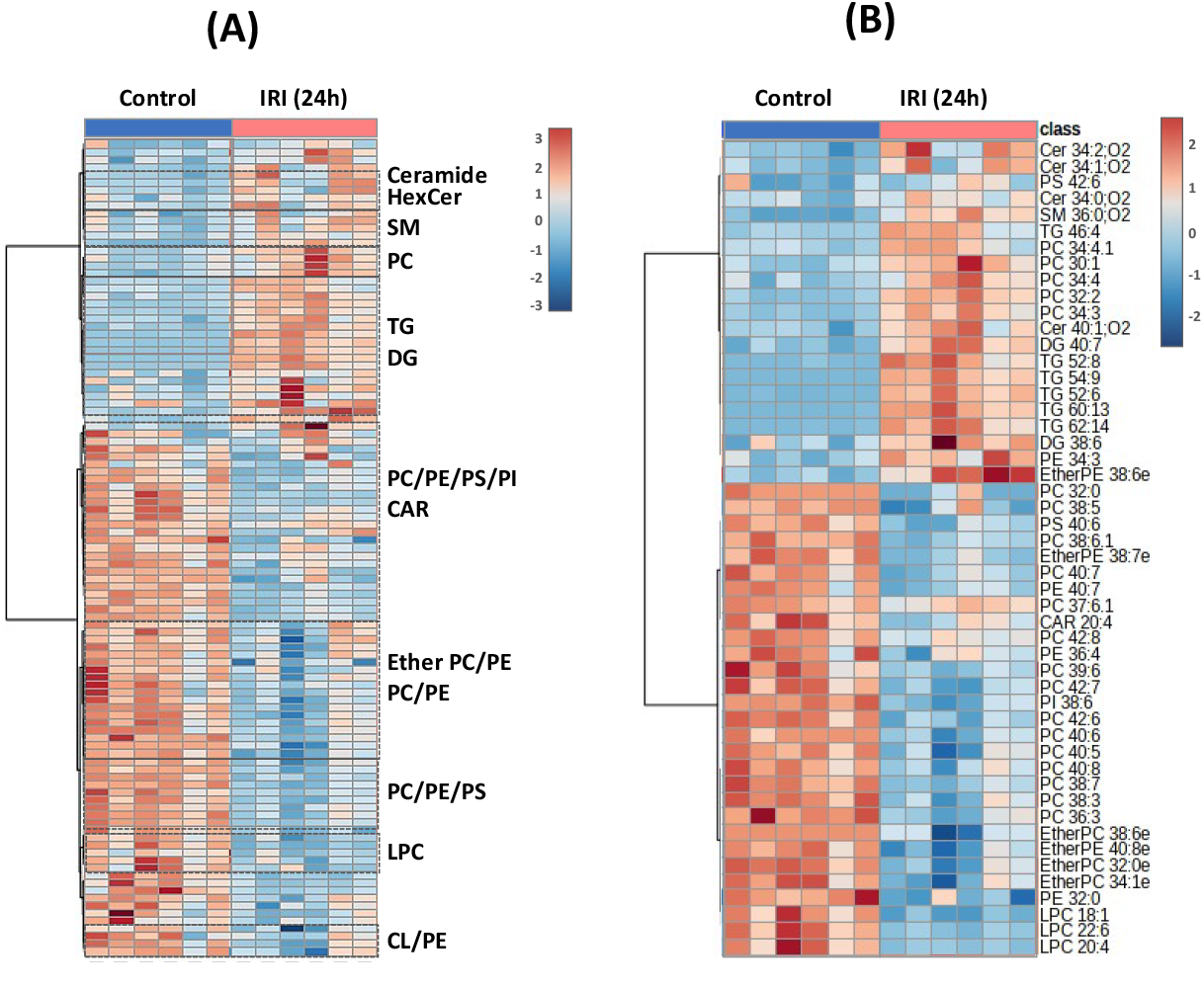
Heatmaps of lipid levels showing significantly altered abundance in kidneys following renal ischemia-reperfusion injury (IRI) compared to mice without kidney injury (Control). A) Heatmaps showing all differential lipid species indicating their class, B) The top 50 differential lipid species. Heatmaps were generated with the Metaboanalyst 6.0 platform (https://www.metaboanalyst.ca), with hierarchical clustering based on Euclidean distance and Ward’s clustering. CAR: Fatty acyl carnitines; Cer: Ceramides; CL: Cardiolipins; DG: Diacylglycerols/Diglycerides; Ether PC: Ether-linked glycerophosphocholines; Ether PE: Ether-linked glycerophosphoethanolamines; HexCer: Hexosylceramides; LPC: Monoacylglycerophosphocholines; PC: Diacylglycerol phosphocholines; PE: Diacylglycerophosphoethanolamines; PI: Diacylglycerol phosphoinositols; PS: Diacylglycerophosphoserines; SM: Sphingomyelins; and TG: Triacylglycerols/Triglycerides.

### Dysregulation of key lipid metabolism genes is an early event in the acute phase of kidney IRI

Changes in lipid levels reflect a dynamic balance between lipid synthesis, lipolysis, and cellular uptake. To identify early changes in lipid-metabolism enzymes that may underlie the lipid changes observed at 24h IRI, we reanalyzed kidney transcriptomic data at 4 h IRI from our previous study [26]. Functional enrichment analysis revealed an early and coordinated transcriptional reprogramming of lipid-metabolism pathways, including phospholipid biosynthesis and metabolism, glycerolipid metabolism, and glycerophospholipid metabolism (**Supplementary Table 2**), suggesting that lipid metabolic dysregulation could be an early feature of kidney IRI.

Clustering analysis of lipid-related differentially expressed genes further highlighted early alterations in FAO enzymes, together with changes in PI, CL, sphingolipid, glycerolipid, and glycerophospholipid metabolism (**Supplementary Figure 2**), suggesting that these early transcriptional responses may shape the lipidomic profile detected at 24h. **Figure 2** integrates these early transcriptomic changes with the lipidomic changes observed at 24h IRI. Quantitative PCR validation confirmed that several of these lipid-metabolism genes also exhibit altered expression at 24h (**Figure 3**), supporting that many lipid changes are preceded by early transcriptional regulation.

**Figure 2.**
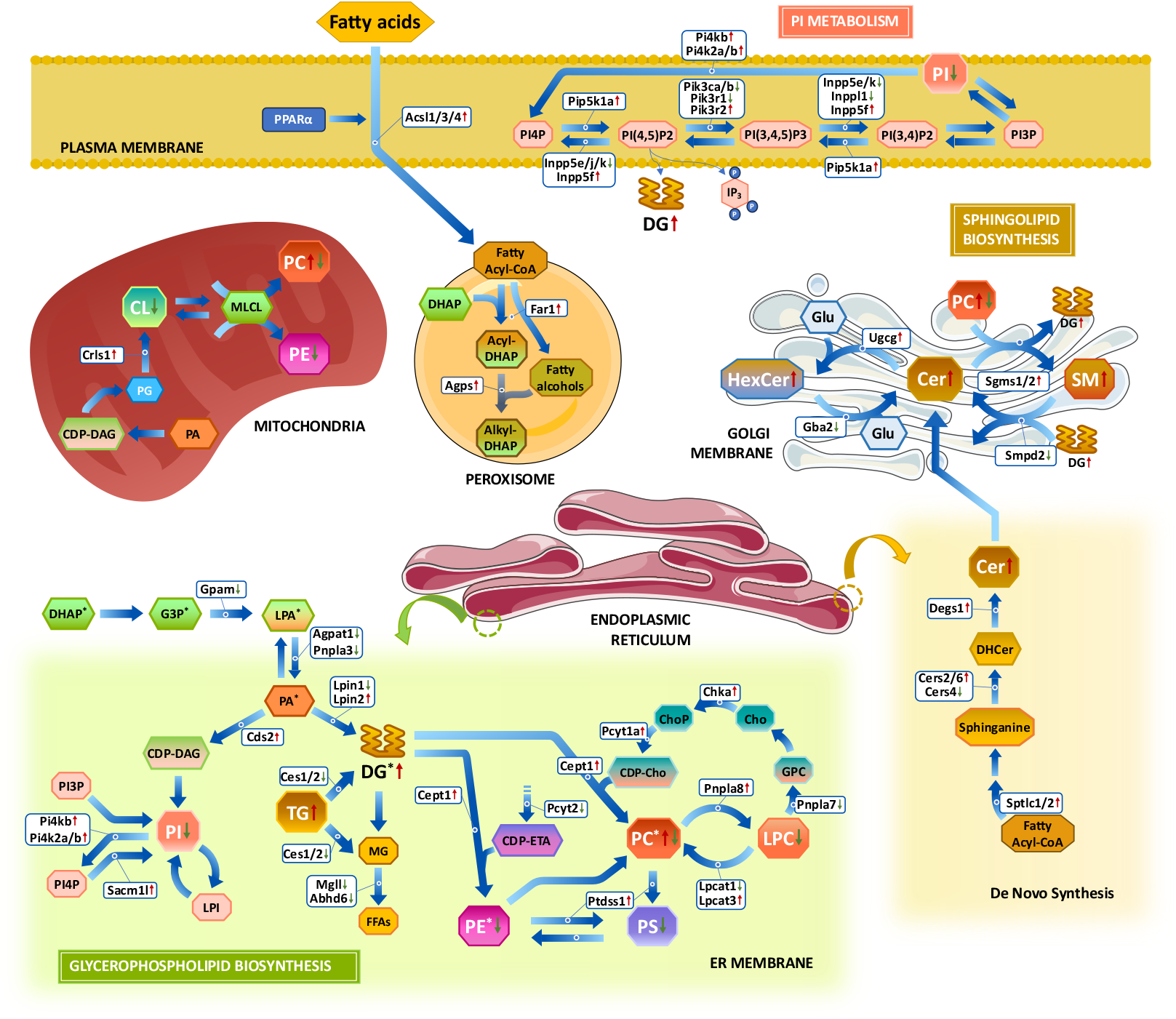
Integrated overview of differentially expressed genes from RNA-seq data of mouse kidneys 4 hours after ischemia–reperfusion injury (IRI_4h) versus sham controls (Ctrl_4h), as reported by Valentijn et al. 2021 [26], and their correspondence with lipidomic alterations observed 24 hours after IRI. The schematic represents phosphatidylinositol (PI) metabolism; the synthesis of diacylglycerols (DG) and triglycerides (TG); glycerophospholipid biosynthesis, cardiolipin (CL) biosynthesis, and sphingolipid biosynthesis. The diagram illustrates the metabolic connectivity between acyl-CoA formation, glycerolipid synthesis, phospholipid turnover, PI signaling, and sphingolipid metabolism, supporting coordinated lipid metabolic reprogramming following kidney ischemic injury. In the endoplasmic reticulum (ER) membrane, the expression of several enzymes implicated in glycerolipid and glycerophospholipid metabolism was found to be altered in early after IRI (IRI-4h). These changes include enzymes involved in phosphatidylcholine (PC) and phosphatidylethanolamine (PE) metabolism, particularly in the Kennedy pathways (CDP-choline and CDP-ethanolamine branches) and the Lands’ cycle. Ether lipid synthesis begins with the formation of alkyl-DHAP in peroxisomes and continues with subsequent reactions in the ER membrane; enzymes involved in ether lipid synthesis are marked with an asterisk (*). Increases and decreases in gene expression at 4h and lipid abundance at 24h following IRI are indicated by red (↑) and green (↓) arrows, respectively. Cer: Ceramide; CL: Cardiolipin; CoA: Coenzyme A; CDP-DAG: CDP-diacylglycerol; DG (DAG): Diacylglycerol; DHCer: Dihydroceramide; DHAP: Dihydroxyacetone phosphate; EtherPC: Ether-linked phosphatidylcholine; EtherPE: Ether-linked phosphatidylethanolamine; FA: Fatty acids; G3P: Glycerol-3-phosphate; Glu: Glucose; GPC: Glycerophosphocholine; HexCer: Hexosylceramide; LPA: Lysophosphatidic acid; LPC: Lysophosphatidylcholine; LPE: Lysophosphatidylethanolamine; MG: Monoacylglycerol; PA: Phosphatidic acid; PC: Phosphatidylcholine; PE: Phosphatidylethanolamine; PG: Phosphatidylglycerol; PI: Phosphatidylinositol; PI3P / PI4P / PI(4,5)P_2_ / PI(3,4,5)P_3_ / PI(3,5)P_2_: Phosphoinositides; PS: Phosphatidylserine; SM: Sphingomyelin; and TG: Triglyceride.

**Figure 3.**
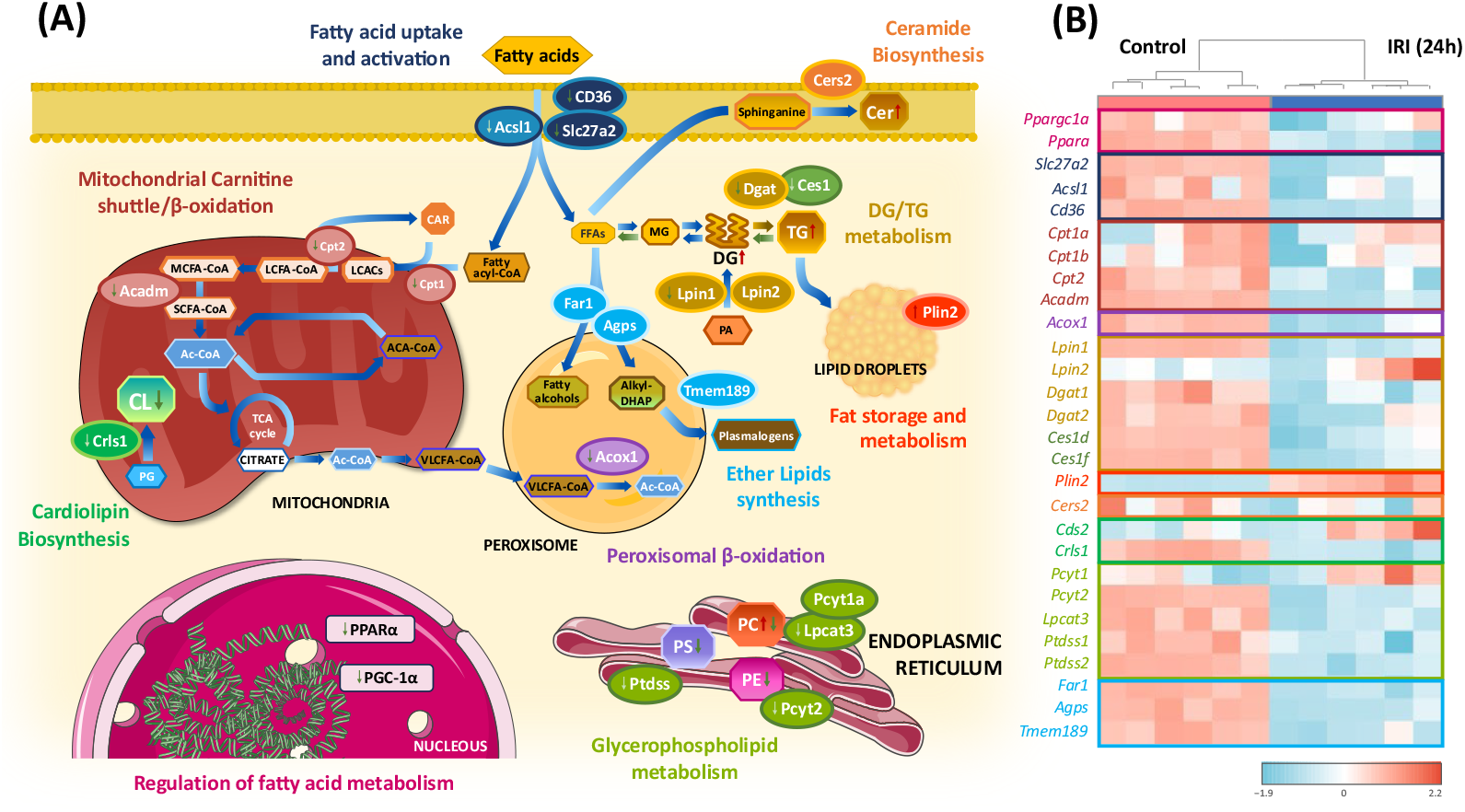
Integrated overview of lipid metabolic reprogramming in the kidney 24h following ischemia–reperfusion injury (IRI). (A) Schematic overview of the main lipid metabolic pathways altered 24h following IRI, including transcriptional regulation of fatty acid metabolism, fatty acid uptake and activation, mitochondrial carnitine shuttle, mitochondrial and peroxisomal β-oxidation, diacylglycerol (DG) and triglyceride (TG) metabolism, lipid droplet formation, ceramide, cardiolipin (CL) and Ether lipid biosynthesis, and glycerophospholipid metabolism. (B) Heatmap showing expression patterns of selected lipid metabolism–related genes validated by qPCR in control and IRI kidneys at 24h, clustered by functional pathways. Red and blue indicate relative up- and down-regulation, respectively. Gene expression data are presented as fold change relative to control (mean ± SEM; n = 6 kidneys per group). Log_2_ fold-change values and statistical significance for each validated gene, together with corresponding transcriptomic data at 4 h post-IRI, are provided in Supplementary Table 2. Acadm: Medium-chain specific acyl-CoA dehydrogenase; Ac-CoA: Acetyl coenzyme A; Acox1: peroxisomal acyl-coenzyme A oxidase 1; Acsl1: Long-chain-fatty-acid--CoA ligase 1; CD36: Platelet glycoprotein 4; Ces1: Carboxylesterase 1; Cpt1: Carnitine O-palmitoyltransferase 1; Cpt2: Carnitine O-palmitoyltransferase 2; Dgat: Diacylglycerol O-acyltransferase; FFAs: Free Fatty Acids; GLPs: Glycerophospholipids LCACs: Long-chain acylcarnitines; LCFA-CoA: Long-chain fatty acyl-CoA; MCFA-CoA: Medium-chain fatty acyl-CoA; MGs: Monoglycerols or monoglycerides; PA: Phosphatidic acid; PI: Phosphatidylinositol; PPARGC1A: Peroxisome proliferator-activated receptor gamma coactivator 1-alpha; PPARα: Peroxisome proliferator-activated receptor alpha; SCFA-CoA: Short-chain fatty acyl-CoA; Slc27a2: Long-chain fatty acid transport protein 2; and VLCFA-CoA: Very long-chain fatty acyl-CoA.

At 4h post-IRI, inositol phosphate metabolism emerged as one of the most significantly enriched lipid-related pathways (**Supplementary Table 2 and Supplementary Figure 2**). Multiple PI kinases, phosphatases, and phosphoinositide-specific phospholipase C (PLC) enzymes were differentially expressed, consistent with early activation of PI turnover and signaling. This transcriptional profile provides a mechanistic basis for the lipidomic pattern observed at 24h, characterized by depletion of PI species and accumulation of DG species (**Figure 2**). In parallel, CDP-diacylglycerol synthase (*Cds2*), which catalyzes the conversion of phosphatidic acid (PA) to CDP-diacylglycerol (CDP-DAG) in the endoplasmic reticulum (ER), was upregulated at 4h but showed only a nonsignificant trend at 24h (**Figures 2 and 3**). Cardiolipin synthase (*Crls1*) followed a similar temporal pattern, with early upregulation and significant downregulation at 24h (p-value=0.0022), consistent with the reduction in cardiolipin species detected at this time point.

Sphingolipid metabolism was also significantly altered early after IRI (**Supplementary Figure 2**). Transcriptomic analysis revealed increased expression of enzymes involved in de novo ceramide synthesis in the ER, together with enzymes mediating ceramide conversion in the Golgi (**Figure 2**). Early upregulation of sphingomyelin synthases (*Sgms1* and *Sgms2*) and ceramide glucosyltransferase (*Ugcg*) supports the subsequent increase in SM and HexCer levels at 24h, respectively. Since *Sgms* enzymes catalyze the conversion of PC and ceramide into SM and DG, these changes provide a direct mechanistic link between sphingolipid metabolism and phospholipid remodeling. The downregulation of sphingomyelin phosphodiesterase 2 (*Smpd2*) and non-lysosomal glucosylceramidase (*Gba2*) further favors SM and HexCer accumulation. Despite downstream accumulation of SM and HexCer, ceramide levels remained elevated at 24h, suggesting that sustained ceramide abundance reflects early activation of de novo synthesis rather than continued transcriptional upregulation at later stages. Consistently, no significant changes were observed at 24h in the expression of *Cers2*, a ceramide synthase highly expressed in the kidney and responsible for very-long-chain ceramide production (**Figure 3 and Supplementary Table 2**) [29].

Changes in multiple genes involved in glycerolipid metabolism were also detected at 4h and 24h (**Figures 2 and 3**). Significant changes were detected in phosphatidate phosphatase (*Lpin1* and *Lpin2*), key magnesium-dependent enzymes that catalyze the dephosphorylation of PA to form DG, an intermediate in TG, PC, and PE biosynthesis. Upregulation of *Lpin2 at 4h*, together with a nonsignificant trend toward increased expression at 24h, was consistent with DG accumulation and subsequent glycerolipid and glycerophospholipid synthesis, while *Lpin1* was downregulated, pointing to isoform-specific functions in hypoxia-driven lipid remodeling (**Figures 2 and 3**). The early downregulation of monoglyceride lipases (*Mgll* and *Abhd6*) and carboxylesterases (*Ces1d, Ces1e, Ces2g*, and *Ces2h*), together with marked suppression of *Ces1d* and *Ces1f* at 24h, also provides a plausible explanation for TG accumulation after IRI through decreased catabolism (Figure 3 and **Supplementary Table 2)**. Decreased degradation combined with increased expression of diacylglycerol O-acyltransferases (*Dgat1* and *Dgat2*), which catalyze the committed step of TG synthesis from DG and fatty acyl-CoA substrates [30], could account for the abrupt rise in TG levels observed after IRI. Nevertheless, the reduced expression of *Dgat* enzymes at 24h suggests that impaired lipid hydrolysis is a major contributor to TG accumulation, although enhanced TG synthesis may represent an earlier, transient event following IRI. Consistent with this hypothesis, TG accumulation was preceded by the early upregulation of the lipid droplet–associated gene perilipin-2 (*Plin2*), which remained elevated at 24h (**Figure 3 and Supplementary Table 2)**.

Multiple enzymes regulating glycerophospholipid biosynthesis were altered early after IRI, particularly within the Kennedy pathways of PC (CDP-choline branch) and PE synthesis (CDP-ethanolamine branch), the Lands cycle of phospholipid remodeling, and ether lipid synthesis, indicating early remodeling of PC, PE, PS, LPC, and ether phospholipids (**Figure 2**). Early upregulation of choline kinase (*Chka*) and choline-phosphate cytidylyltransferase A (*Pcyt1a*) supports enhanced PC biosynthesis. In parallel, differential regulation of lysophosphatidylcholine acyltransferases, including the downregulation of *Lpcat1* and upregulation of *Lpcat3*, indicates a shift toward PC remodeling via the Lands cycle. At 24h, however, *Lpcat3* expression was reduced and *Pcyt1a* unchanged, indicating attenuation of both LPC reacylation and de novo PC synthesis (**Figure 3 and Supplementary Table 2**). This temporal transition explains the coexistence of increased and decreased PC species at 24h, reflecting early synthesis and remodeling followed by selective turnover, while persistently low LPC levels are consistent with efficient early reacylation and limited LPC regeneration. Conversely, key components of the CDP-ethanolamine pathway, including *Pcyt2*, were downregulated at both 4h and 24h, providing a mechanistic explanation for the reduction in PE species (**Figures 2 and 3**). Increased expression of choline/ethanolaminephosphotransferase 1 (*Cept1*) at 4h further supports phospholipid remodeling. In addition, phosphatidylserine synthases *Ptdss1* and *Ptdss2* were significantly reduced at 24h, reinforcing the decrease in PS species. We also observed an early upregulation of fatty acyl-CoA reductase 1 (*Far1*) and alkyldihydroxyacetonephosphate synthase (*Agps*), enzymes responsible for ether lipid synthesis [31], which suggests a transient compensatory activation of this peroxisomal pathway (**Figure 2**). However, downregulation of these enzymes at later time points, together with reduced expression of downstream ER enzymes and *Tmem189*, aligns with the decreased EtherPC and EtherPE species observed at 24h, indicating that early compensatory activation of ether-lipid synthesis is not sustained during ongoing injury (**Figures 2 and 3**).

In addition to pathway-specific alterations in lipid synthesis, we evaluated broader disruptions in renal lipid homeostasis arising from imbalances in fatty acid uptake and activation, β-oxidation, lipid storage, and lipolysis by quantitative PCR (**Figure 3 and Supplementary Table 2)**. Key regulators of lipid metabolism, including peroxisome proliferator-activated receptor alpha (PPARα) and its coactivator PGC-1α (Ppargc1a), were significantly suppressed at 24h after IRI, whereas *Ppargc1a* was already reduced at 4h (**Figure 3**). Consistently, crucial FAO enzymes, including carnitine O-palmitoyltransferases (*Cpt1* and *Cpt2*), mitochondrial medium-chain specific acyl-CoA dehydrogenase (*Acadm*), and peroxisomal acyl-coenzyme A oxidase 1 (*Acox1*), were downregulated at 24h (**Figure 3 and Supplementary Table 2**). Transcriptomic reanalysis further revealed that several peroxisomal FAO enzymes were already suppressed at 4h post-IRI (**Supplementary Table 2**). Our results also suggest an early reduction in cellular lipid uptake and activation. While long-chain fatty acid transport protein 2 (*Slc27a2*), the fatty acid translocase *CD36* (platelet glycoprotein 4), and the long-chain-fatty-acid--CoA ligase 1 (*Acsl1*) were significantly downregulated at 24h, several other fatty acid transporters and lipid-activating enzymes were already decreased at 4h following IRI (**Figure 3 and Supplementary Table 2**).

### Insights from single-cell transcriptomics

Single-cell transcriptomics datamining using publicly available KIT datasets [28,32] allowed the mapping of lipid metabolism–related transcriptional changes to specific renal cell populations, highlighting proximal tubular cells as a major contributor to the lipid abnormalities observed after IRI (**Figure 4A**).

**Figure 4.**
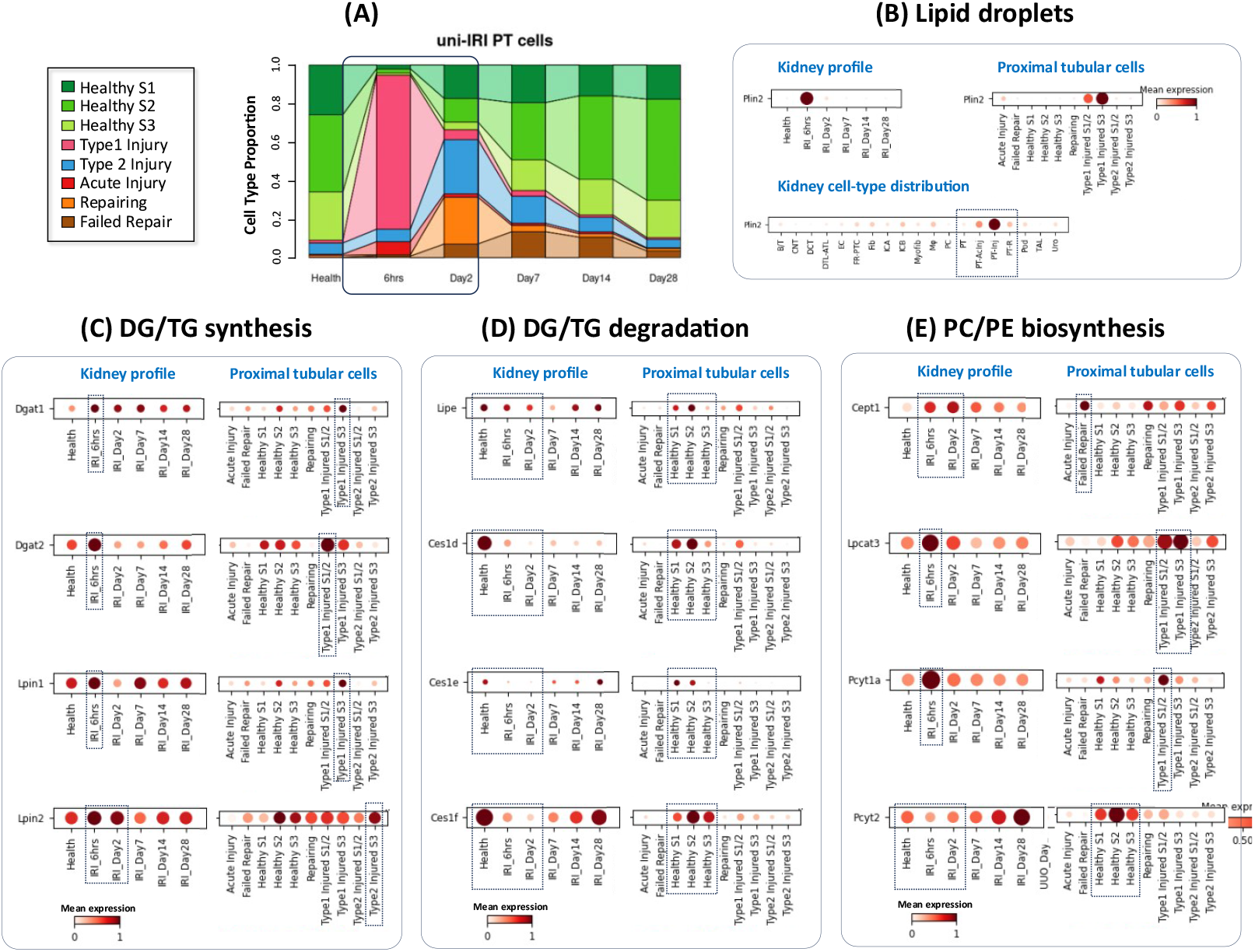
Single-cell transcriptomic data mining using Kidney Interactive Transcriptomics (KIT) datasets (http://humphreyslab.com/SingleCell/) [28,32] to provide a more complete overview of molecular mechanisms in kidney and proximal tubular (PT) cells involved in lipid abnormalities. Dot plots show mean gene expression (color intensity) and proportion of expressing cells (dot size). A) Different proximal tubular cell types based on single-cell transcriptomics profiles following murine acute ischemia–reperfusion injury (IRI-AKI). Type 1 injured PT cells are most abundant within the first 6 h following IRI, accounting for approximately 80% of PT cells, and represent a transient injury state with recovery potential. In contrast, Type 2 injured PT cells are more characteristic of maladaptive injury contexts and could progress toward a failed-repair proximal tubular cell (FR-PTC) state that promotes fibrogenesis. B) Perilipin-2 (Plin2), a lipid droplet– associated gene, is selectively overexpressed in type 1 injured proximal tubular cells. C) Mean expression of genes encoding enzymes involved in diacylglycerol (DG) and triglyceride (TG) synthesis demonstrates early induction of Diacylglycerol O-acyltransferase (Dgat) isoforms and phosphatidate phosphatase Lpin1 predominantly in Type 1 injured PT cells, followed by downregulation by 48h. In contrast, Lpin2 expression is more closely associated with Type 2 injured PT cells. (D) Mean expression of genes encoding enzymes involved in TG degradation. Multiple enzymes display a decreased expression, starting at 6 h and persisting over time. The decrease is shared by type 1 and type 2 injured proximal tubular cells, the latter predominant at later timepoints following IRI, as well as by persistently injured failed repair cells. (E) Mean expression of genes encoding enzymes implicated in the phosphatidylcholine (PC) and phosphatidylethanolamine (PE) biosynthesis. Cept1 is associated with failed repair in tubular cells. Lpcat3 and Pcyt1a enzymes are associated with remodeling via the Kennedy pathway and Lands’ cycle, are transiently induced early after IRI and are preferentially expressed in Type 1 injured PT cells, with reduced expression in failed-repair PT cells. In contrast, Pcyt2, a key enzyme of the CDP-ethanolamine pathway for PE synthesis, is enriched in healthy PT cells and downregulated in injured PT states.

The pattern of high TG and DG levels observed at 24h could be traced to transient type 1 injured proximal tubular cells, characterized by the expression of the lipid droplet-related gene *Plin2* [28] (**Figure 4B**). These cells display coexistent early upregulation of genes encoding glycerolipid synthesis enzymes such as *Dgat* and *Lpin* and downregulation of genes encoding triglyceride degradation enzymes upon IRI (**Figure 4C/D**). A more persistent decrease in TG degradation can be traced to the downregulation of multiple genes encoding lipolysis enzymes, which is observed in all injured proximal tubular types, including those with failed repair that fail to return to a physiological phenotype. In summary, TG accumulation appears to result from coordinated gene expression programs involving several cell subtypes and multiple genes regulating both TG synthesis and degradation.

Single-cell transcriptomic data mining further provided insight into the cellular origin of PC and PE remodeling observed in our lipidomic analyses (**Figure 4E**). *Cept1*, which catalyzes the final step of PC and PE synthesis in the Kennedy pathway, was early induced in proximal tubular cells and was expressed in both Type 1 and Type 2 injured proximal tubular cells populations. This induction persisted during early repair, peaking at 48 h post-IRI, mainly in failed-repair proximal tubular cells, suggesting that early activation of the Kennedy pathway may contribute to maladaptive repair. A transient induction of PC *biosynthetic* and remodeling enzymes, including *Pcyt1a* (CDP-choline/Kennedy pathway) and *Lpcat3* (Lands’ cycle), was observed at 6 h post-IRI. Data mining traced this response predominantly to Type 1 injured proximal tubular cells, indicating an early, adaptive upregulation of PC biosynthesis and remodeling during the acute injury phase, followed by attenuation at later time points. This temporal pattern reinforces our integrated transcriptomic and lipidomic findings. In contrast, *Pcyt2*, a key enzyme of the CDP-ethanolamine pathway responsible for PE biosynthesis [33], was preferentially expressed in healthy proximal tubular cells and downregulated early after injury, suggesting that PE synthesis is linked to a more protective, homeostatic phenotype. Collectively, these findings indicate that PC remodeling could primarily be driven by transiently injured proximal tubular cells, particularly Type 1 injury cells, whereas PE biosynthesis is associated with non-injured cells and with recovery potential. This cell–type–specific transcriptional organization provides a mechanistic framework linking early proximal tubular injury states to the divergent lipidomic signatures observed in IRI-AKI.

### Lipid species and lipid metabolism enzymes correlate with markers of kidney injury and nephroprotective factors in IRI-AKI

To explore the biological and functional significance of lipidomic alterations, we evaluated potential correlations between specific lipid species and genes involved in lipid metabolism, biomarkers of kidney damage, proinflammatory, redox-related, and nephroprotective factors changed following IRI (**Figure 5A**), as well as renal function parameters (**Supplementary Figure 3**). Data mining with the KIT platform showed that several of these genes have already been implicated in metabolic dysfunction and tubular injury in IRI-AKI [28] (**Supplementary Figure 4**). Global correlation analysis revealed distinct metabolic and molecular modules associated with either kidney damage or nephroprotection following IRI (**Figure 5B, Supplementary Table 3, and Supplementary Figures 5-8**).

**Figure 5.**
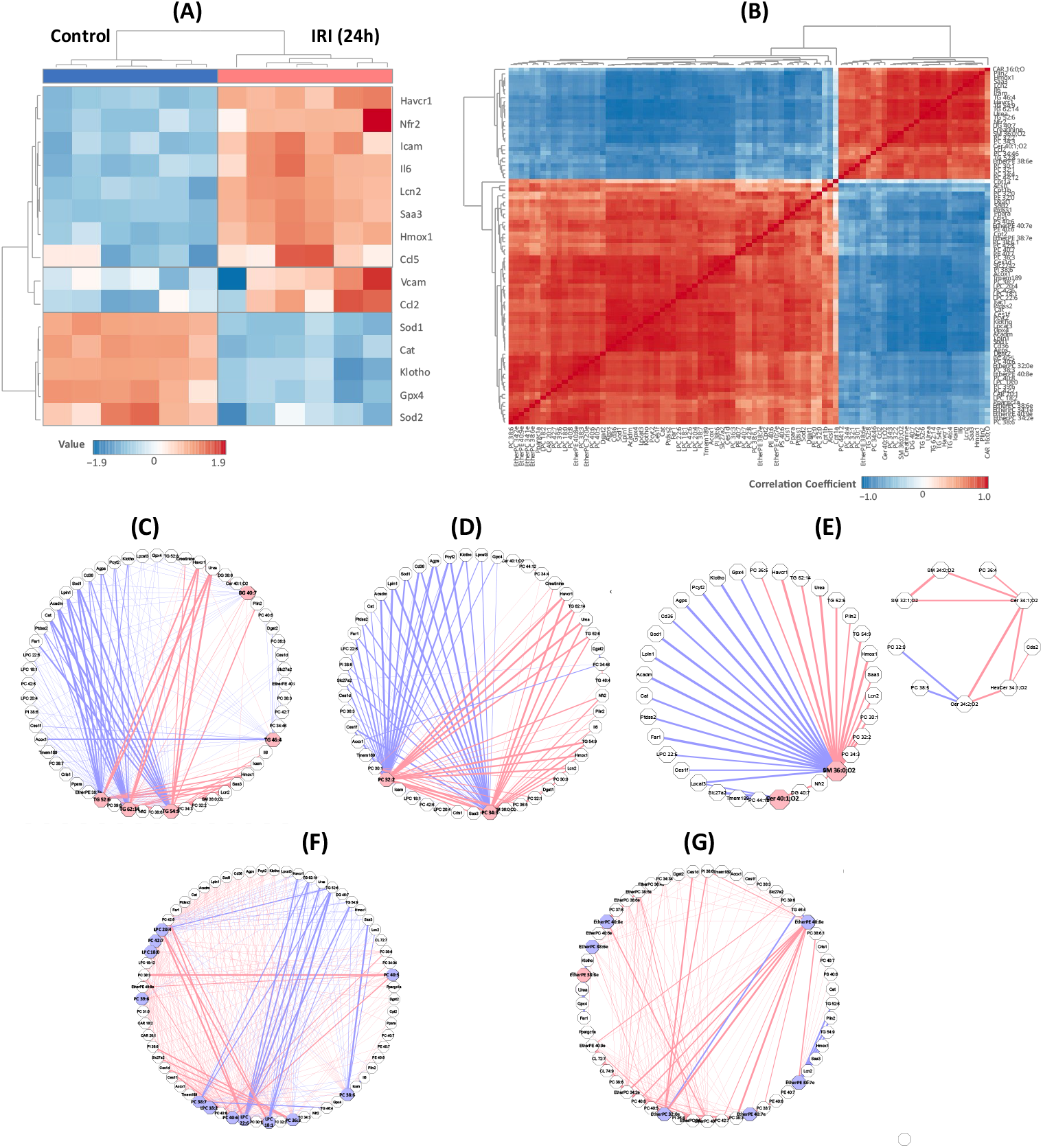
Correlation analysis between lipid species and the expression of lipid metabolism enzymes, tubular injury biomarkers, inflammatory mediators, nephroprotective factors such as Klotho, and antioxidant enzymes, as well as classical renal function markers (urea and creatinine). (A) Heatmap showing the expression of key renal injury, inflammatory, antioxidant, and nephroprotective genes in kidneys from ischemia–reperfusion–injured (IRI) and control mice. Genes include tubular injury markers (Havcr1, Lcn2), inflammatory mediators (Il6, Ccl2, Vcam1), antioxidant enzymes (Sod1, Cat, Gpx4, Hmox1), and the nephroprotective factor Klotho. (B) Correlation matrix showing correlation of the top 50 lipids with qPCR genes and renal function markers. Correlation was based on the Pearson correlation, and the coefficients revealed distinct lipid–gene co-regulation modules associated with kidney damage or protection. Positive correlations are indicated by red, while negative correlations are indicated by blue. (C–G) Correlation network maps displaying lipid– gene interactions for major lipid classes, including (C) triglycerides (TGs) and diglycerides (DGs); (D) phosphatidylcholines (PCs); (E) sphingomyelins (SMs) and ceramides (Cer); (F) phosphatidylcholines (PCs) and lysophosphatidylcholines (LPCs); and (G) ether-linked phospholipids (EtherPC and EtherPE). Networks were generated using Metscape (Cytoscape app) based on Pearson correlation coefficients higher than 0.8. (adjusted p < 0.05). Edge thickness reflects correlation strength, with pink edges indicating positive and purple edges negative associations. Panels (C–E) represent lipid species that increased following injury, whereas panels (F–G) depict lipids that decreased after IRI and correlated with nephroprotective or antioxidant profiles. The most relevant lipids that are positively correlated with damaging factors are highlighted in pink, while those that are positively correlated with nephroprotectors are highlighted in purple. **Abbreviations:** Cat: Catalase; Ccl2: C-C motif chemokine 2;Ccl5: C-C motif chemokine 5; GPX4: Glutathione Peroxidase 4; Havcr1: Hepatitis A virus cellular receptor 1; Hmox1: Heme oxygenase 1; Icam-1: Intercellular adhesion molecule 1; Il6: Interleukin-6; Lcn2: Neutrophil gelatinase-associated lipocalin; Nfe2l2: Nuclear factor erythroid 2–related factor 2; Sod: Superoxide dismutase; and Vcam: Vascular cell adhesion protein.

Among lipids accumulated after IRI, several glycerolipids (TG and DG), sphingolipids (e.g., SM 36:0;O2), and PCs showed strong positive correlations (r >0.8) with classical markers of renal dysfunction (urea, creatinine), tubular injury (*Havcr1*, encoding for KIM-1), and renal damage (*Lcn2*, encoding for NGAL) (**Figures 5C-E and Supplementary Figure 6**). The most notable correlations (r >0.9) were observed for TG 54:9, TG 52:6, TG 62:14, DG 40:7, PC 32:2, and PC 34:3 (**Supplementary Figure 6**), with TG 54:9 being detected exclusively after ischemic injury, underscoring its potential as a specific biomarker of renal damage. DG 40:7 and several TGs (TG 62:14, TG 54:9, TG 52:6) and PCs (PC 34:3, PC 32:2) correlated positively with the oxidative-stress sensors *Nrf2* and *Hmox1* (**Figures 5C/D)**, which are also overexpressed in type 1 injured proximal tubular cells (**Supplementary Figures 4/8**), and upregulated after IRI, reflecting an adaptive antioxidant response.

Inflammatory mediators exhibited similar trends, displaying robust correlations with TGs, particularly TG 46:4, which correlated strongly (r >0.8) with several pro-inflammatory markers, including *Il6, Ccl2*, and *Vcam1 (***Figure 5C/D and Supplementary Figure 7**). Several PCs (notably PC 34:3 and PC 32:2) also correlated strongly (r >0.9) with acute-phase (*Saa3*), cytokine (*Il6*), and adhesion-related (*Icam1*) genes, supporting the role of specific PC species in inflammation-driven injury (**Figure 5D and Supplementary Figure 7**). Interestingly, *Ccl2* and *Vcam1* exhibited distinct correlation patterns. While *Ccl2* showed a strong association with TG 46:4 and the very-long-chain ceramide 40:1;O2 (but not with the fully saturated Ceramide 40:0;O2), *Vcam1* showed only modest correlations (**Supplementary Figure 7**). Again, this may be understood using single-cell transcriptomics datamining, as *Vcam1* is upregulated in failed repair proximal tubular cells. Failed repair cells derive mainly from type 2 injured cells, which differ from the type 1 injured cells that upregulate the molecular machinery to accumulate TG [28] (**Supplementary Figure 4**). Together, these patterns suggest that the accumulation of specific TGs/DGs, PCs, and ceramides during AKI is associated with more severe tubular injury, enhanced inflammatory activation, and poorer renal outcomes, highlighting their potential as lipid-based biomarkers of kidney damage.

On the other hand, these damage-associated lipids correlated negatively with antioxidant enzymes (e.g., *Sod1, Cat*), FAO enzymes (e.g., *Acadm*), lipid transporters (e.g., *Cd36*), and lipid-metabolism enzymes (e.g., *Pcyt2, Ces1*), while correlating positively with the lipid-droplet protein Plin2. As shown above, Plin2 marks transient type 1 injured proximal tubular cells, which are characterized by TG and DG accumulation following IRI (**Figure 4B**). In contrast, antioxidant and lipid-metabolic enzymes, which were correlated here with nephroprotective markers, such as *klotho* and glutathione peroxidase 4 (*Gpx4*), are predominantly associated with healthy or recovery-prone tubular phenotypes in single-cell transcriptomic analyses [28] (**Figure 4 and Supplementary Figure 9**). Together, these associations suggest that accumulation of these damage-associated lipid species reflects a metabolic blockade in injured proximal tubular cells, occurring within an oxidative-stress–permissive environment that favors lipid storage over oxidation and turnover. Also reinforcing this, the mitochondrial antioxidant Sod2 showed the strongest negative correlation with CAR 16:0;O, the only acylcarnitine that increased following IRI (**Supplementary Figure 10**). CAR 16:0;O also correlated negatively with Dgat enzymes and positively with damage-associated lipids such as SM 36:0;O_2_, underscoring its potential as a biomarker of impaired lipid metabolism and mitochondrial function, and oxidative stress.

Conversely, a subset of phospholipids displayed an association with a nephroprotective phenotype, including PE 40:6 and PI 38:6, as well as several PC/LPC and ether-linked phospholipids. These lipid species correlated positively with nephroprotective markers (*Klotho, Gpx4*), as well as with antioxidant enzymes (*cat, sod1*), and inversely with markers of renal dysfunction, tubular injury, and inflammation (**Figure 5 F/G and Supplementary Figures 6-8**). This correlation pattern is associated with a healthier tubular phenotype, as supported by single-cell transcriptomic data (**Supplementary Figure 4**). The strongest correlations (r >0.8) were established between *Klotho*/*Gpx4* and PI 38:6 and several LPCs (LPC 20:4, LPC 18:1, LPC 22:6) and specific PCs (PC 36:3, PC 38:7, PC 40:6) (**Supplementary Figure 6**). These nephroprotection-associated PCs and LPCs also showed very strong negative correlations with serum creatinine, injury markers, and damage-associated triglycerides, supporting their association with preserved renal function and a protective metabolic profile (**Figure 5F and Supplementary Figure 6**). In contrast, damage-associated PCs (e.g., PC 34:3, PC 32:2) exhibited an opposite remodeling trend, correlating positively with TG accumulation and inflammatory mediators, indicative of an active but maladaptive phospholipid remodeling during injury progression.

Another lipid class that exhibited a consistent association with a nephroprotective pattern was ether-linked phospholipids (EtherPC, EtherPE). Among these species, EtherPC 38:6e and EtherPC 32:0e showed the strongest positive correlations with Klotho (r >0.9; **Figure 5G**), supporting their association with a protective metabolic phenotype. Furthermore, EtherPE 38:7e showed one of the strongest positive correlations with antioxidant *cat*, as well as inverse correlations with *Lcn2, Saa3*, and TGs (TG 54:9, TG 52:6), consistent with an antioxidant and nephroprotective profile (**Figure 5G**). In contrast, EtherPE 38:6e displayed an opposite trend, correlating positively with damage markers and negatively with Klotho, which may reflect a compensatory response to oxidative stress.

Furthermore, these nephroprotection-associated lipid species correlated with several ferroptosis regulators. Strong positive correlations were observed between specific PCs, LPCs, and ether lipid species and key ferroptosis regulators, including *Gpx4, Lpcat3*, and the *Far1–Agps–Tmem189* axis (**Figures 5F/G and Supplementary Table 3**), a pathway essential for the synthesis of ether lipids, including plasmalogens, and recently identified as a ferroptosis regulator [34,35] (**Figures 5C/D and Supplementary Figure 4C**). As the *Far1–Agps* pathway drives the peroxisomal synthesis of ether lipid precursors, its downregulation after IRI was accompanied by a coordinated loss of ether phospholipids, consistent with impaired ether-lipid biosynthesis and increased vulnerability to lipid peroxidation. Concordantly, single-cell transcriptomics revealed that *Gpx4* and *Far1-Agps–Tmem189* expression was predominantly associated with healthy tubular phenotypes [28], reinforcing a functional link between ether-lipid metabolism and ferroptosis protection (**Supplementary Figures 4 and 9**).

Together, these patterns reveal that lipidomic remodeling tightly reflects the metabolic, inflammatory, and redox status of the kidney, offering potential candidate lipids as biomarkers of either injury severity or recovery potential. Accordingly, we defined two major lipid modules: (i) a damage-associated module, which correlated positively with inflammation, FAO suppression, and oxidative stress, and (ii) a nephroprotection-associated module, which correlated with Klotho, lipid metabolism, and antioxidant enzymes (**Figure 6 and Supplementary Table 3**).

**Figure 6.**
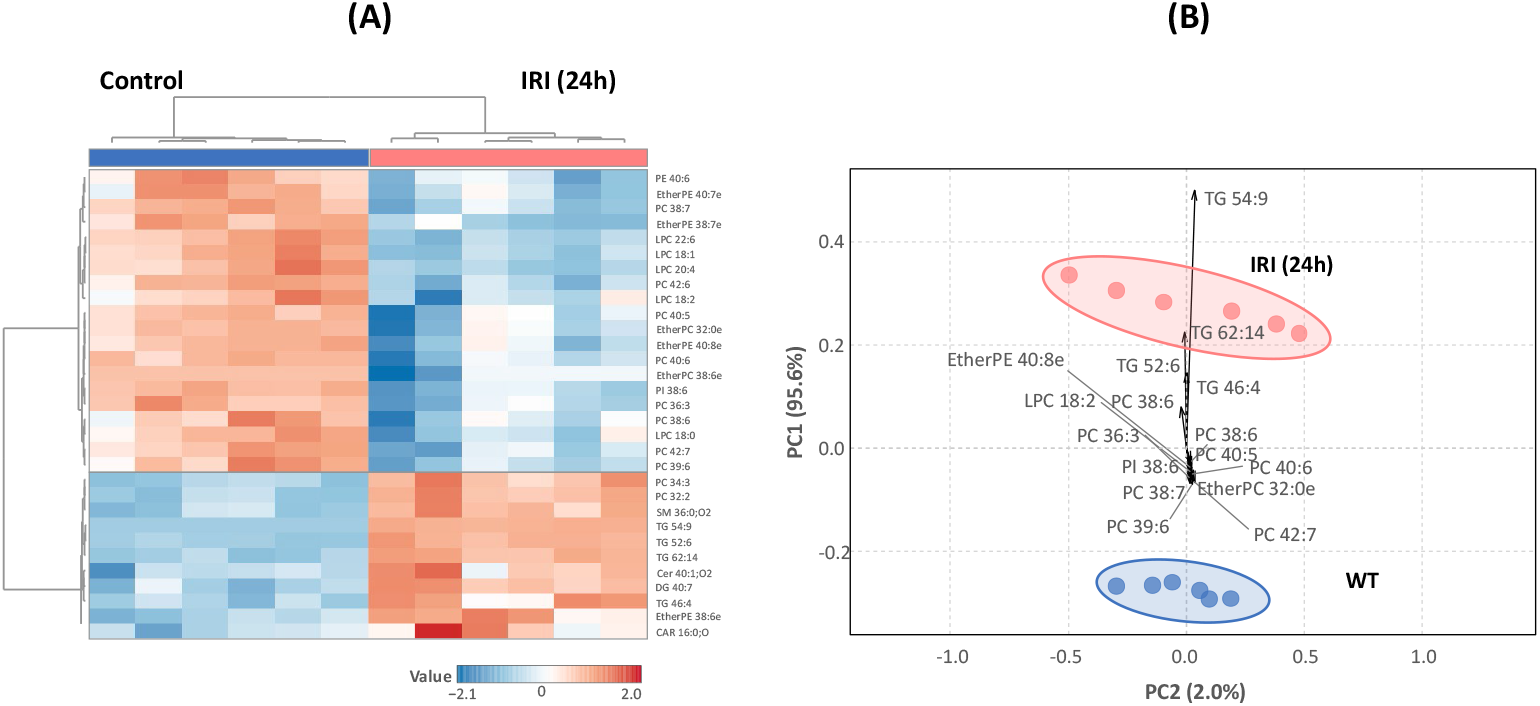
Selected lipid biomarker panel differentiates damage- and protection-associated lipid modules in renal ischemia–reperfusion injury (IRI). (A) Heatmap of the selected lipid biomarker panel showing distinct lipidomic profiles between ischemia–reperfusion injury (IRI) and control kidneys. Hierarchical clustering reveals two major lipid modules: a damage-associated module comprising triglycerides (TGs), diacylglycerols (DGs), saturated phosphatidylcholines (PCs), sphingomyelins/ceramides (SM/Cer), and the acylcarnitine CAR 16:0;O, and a protective module including PCs/PEs, lysophosphatidylcholines (LPCs), phosphatidylinositols (PIs), and ether-linked phospholipids (EtherPC/EtherPE). Red and blue colors indicate increased and decreased abundance, respectively, following IRI. **(B)** Principal component analysis (PCA) based on this biomarker panel, which discriminates between control (blue ellipse) and IRI (red ellipse) kidneys.

## Discussion

This study provides a comprehensive characterization of lipidomic remodeling during the acute phase of kidney IRI, revealing profound alterations in lipid composition that may influence the balance between injury and repair. Our findings extend previous observations of lipid dysregulation in AKI by identifying distinct lipid signatures associated with damage and nephroprotection. Reanalysis of transcriptomic data from our previous work [26] revealed that early transcriptional alterations following IRI precede and likely drive the lipidomic profiles observed at 24h, shifting toward the accumulation of lipotoxic species (TGs, DGs, Cer) and the concomitant loss of protective lipids (Ether Lipids, PE). These findings reinforce the concept that metabolic reprogramming during IRI is an early and decisive determinant of tubular injury and maladaptive repair [10,13,36], impacting not only FAO but also substantially altering lipid biosynthesis and catabolism.

Several authors reported changes in renal lipid levels as a common phenotype following IRI-mediated renal injury [37–39]. Our lipidomic results confirmed that substantial alterations in renal lipid composition are induced following IRI, identifying 84 significantly altered lipid species in IRI-injured kidneys. Early transcriptomic changes observed 4h after IRI suggest a rapid suppression of key regulators of fatty acid metabolism, in particular the PPARα coactivator PGC-1α, which orchestrate mitochondrial biogenesis and β-oxidation [40]. This downregulation, together with the reduced expression of critical FAO enzymes at 24h, including *Acadm* and *Acox1*, and accumulation of specific acetylcarnitines (CAR 16:0;O), likely underlies the impaired lipid catabolism and subsequent lipid accumulation detected at 24h post-IRI. The decline in the expression of key fatty acid transporters (*Slc27a2* and *Cd36*), lipid-activating enzymes (*Acsl1*), and components of the mitochondrial carnitine shuttle (*Cpt1b and Cpt2*), some already evident at 4h post-IRI, indicates an early impairment of mitochondrial fatty acid import. Since carnitine shuttle constitutes a rate-limiting step for mitochondrial FAO, these alterations strongly support compromised FAO capacity during the acute phase of injury [41]. FAO represents the principal energy source for the kidney, particularly for proximal tubular cells, its impairment may trigger an early maladaptive metabolic switch characterized by ATP depletion, mitochondrial dysfunction, and intracellular lipid accumulation, thereby promoting tubular injury [11,42]. In parallel, several peroxisomal FAO enzymes were also downregulated at 4 h post-IRI, indicating that peroxisomal FAO is also affected early after ischemic injury. Given the prominent role of peroxisomal FAO in proximal tubular cells, where Acox enzymes are selectively expressed, these findings are consistent with experimental ischemia models reporting early peroxisomal FAO impairment during tubular injury [43]. Therefore, therapeutic strategies aimed at restoring mitochondrial and peroxisomal FAO, through PPARα agonists, carnitine supplementation, or modulation of CPT activity, may help prevent lipid accumulation and mitigate AKI progression [7].

Progressive accumulation of TGs in tubular epithelial cells is a well-recognized feature of ischemic injury, and our results align with prior work linking ischemia-induced hypoxia to reduced fatty acid transport and inhibition of FAO, thereby diverting lipid flux toward TG storage [39,44–46]. Given the kidney’s reliance on FAO, such metabolic inflexibility is thought to contribute to ATP depletion, mitochondrial dysfunction, inflammatory signaling, and CKD progression [8,10,11,42,47]. In our study, glycerolipids emerged as the most prominently altered lipid class following IRI, with a marked accumulation of highly unsaturated long-chain TGs (TG 52:8, TG 54:9, and TG 60:13) detectable only in injured kidneys. Concomitant increases in DGs and ceramides further support the emergence of a lipotoxic environment. Indeed, this lipid pattern, characterized by increased TG and ceramide levels, together with reduced CL levels, closely mirrors lipidomic signatures described in kidneys from donors with impaired renal function [48], supporting a translational relevance. This is reinforced by our results, with several TGs and DGs in our dataset showing strong positive correlations with renal functional decline, tubular injury markers (*Havcr1* and *Lcn2*), pro-inflammatory mediators, and oxidative-stress sensors (*Nrf2* and *Hmox1*), which have been found expressed in Type 1 injured proximal tubular cells in response to ischemia-induced damage [28]. Among these species, TG 54:9, detected exclusively after ischemic injury, emerged as a particularly robust candidate biomarker of acute tubular damage, supporting its potential utility for early detection of kidney injury.

Tubular cells are among the most energy-demanding cells in the kidney, making them particularly vulnerable to hypoxic injury and key contributors to CKD progression [49]. Our findings integrate with and extend recent single-cell transcriptomic studies that have helped to define the early metabolic states adopted by proximal tubular cells following ischemic injury. These studies describe a transient type-1 injured proximal tubular cell state marked by rapid TG accumulation and marked induction of Plin2, a lipid-droplet–associated protein tightly linked to metabolic remodeling [28]. A sharp rise in renal TG content within 6 hours of IRI, coincided with the emergence of Plin2+ proximal tubular cells that accumulate lipid droplets and transiently activate lipid-metabolic programs, an adaptive response likely aimed at buffering acute metabolic stress. In this transient type 1 injured proximal tubules, early Plin2 induction coincides with upregulation of glycerolipid synthesis genes (*Dgat, Lpin*) and sustained repression of multiple TG-degrading enzymes beginning at 6 hours post-IRI and persisting thereafter [28]. Consistent with these findings, our transcriptomic analysis revealed early *Plin2* upregulation at 4h post-IRI that persisted at 24h, together with early and sustained downregulation of lipolytic pathways, including multiple monoglyceride lipases and carboxylesterases. These convergent datasets support a model in which early activation of glycerolipid synthesis enzymes, lipid droplet biogenesis, and prolonged suppression of TG catabolism collectively impair TG turnover in injured proximal tubular cells. While initially protective by limiting unesterified fatty acid accumulation during energy stress, this program becomes maladaptive, driving persistent lipid accumulation. Importantly, our data identify defective TG turnover, rather than increased lipid supply, emerges as a key metabolic bottleneck influencing tubular repair. Thus, early metabolic reprogramming establishes a TG-rich state that, if unresolved, may hinder mitochondrial recovery and promote maladaptive repair. Therapeutic strategies to restore lipolytic competence or normalizing TG flux may represent promising avenues to enhance metabolic resilience and improve functional recovery following AKI.

In this metabolic framework, elevated TGs in injured proximal tubular cells are often attributed to both increased synthesis and reduced catabolism [36,44,50]. Nevertheless, our data provide limited support for sustained TG biosynthesis, as *Dgat* isoforms were downregulated at 24h post-IRI, indicating that TG synthesis is a transient, short lived response to injury rather than a persistent driver of lipid accumulation. Instead, the coordinated and durable repression of Ces family enzymes highlights defective TG mobilization as the dominant mechanism at later time points, in agreement with previous studies showing that Ces downregulation exacerbates lipid accumulation and worsens AKI outcomes [51]. The context-dependent roles of Dgat enzymes further underscore the complexity of TG dynamics in renal injury. Dgat1 upregulation has been linked to TG accumulation, impaired bioenergetics, and mitochondrial dysfunction in injured tubules [44], yet Dgat1 is also essential for autophagy-dependent lipid droplet biogenesis, a process that protects mitochondria by sequestering cytotoxic lipid intermediates [52]. Thus, while Dgat1 activation may contribute to maladaptive remodeling under certain conditions, its inhibition can be detrimental in nutrient-limited states such as ischemic stress, underscoring the importance of metabolic state when interpreting lipid handling pathways in AKI. Taken together, our findings reinforce a model in which impaired lipolysis, rather than enhanced lipid synthesis, emerges as the primary mechanism sustaining TG accumulation after IRI. This persistent defect in TG turnover likely amplifies metabolic stress, compromises mitochondrial resilience, and promotes lipotoxic injury, highlighting TG catabolism as a potentially actionable therapeutic axis to enhance metabolic resilience and improve tubular recovery after ischemic AKI.

Phospholipids are major components of cell membranes, and their dysregulation can impair membrane integrity and signaling processes [36]. PC and PE are the most abundant membrane phospholipids, and their ratio affects membrane fluidity, energy metabolism, oxidative stress response, and lipid droplet formation [33]. Their dysregulation is implicated in various developmental and pathological processes, including renal injury [33,37,53]. In our dataset, phospholipids were broadly reduced after IRI, although specific species of PC, PE, and EtherPE deviated from this general decline. This heterogeneity suggests dynamic lipid remodeling rather than uniform phospholipid loss, consistent with previous reports [38,54]. A transient increase in ether-linked phospholipids was observed post-IRI in a previous study, with both PC O-38:1 and PE O-42:3 rising at 6h. However, while PC O-38:1 remained elevated at 24h, PE O-42:3 declined, highlighting divergent remodeling dynamics between PC and PE species during injury progression [37]. A similar pattern was reported in plasma lipidomics, where serum PC levels progressively declined across CKD stages, pointing to an early transient increase of these lipid species followed by depletion [55], again supporting the translational relevance of the present findings. Our data provides a mechanistic framework for the altered PC, PE, and LPC patterns observed post-IRI. We identified a transient induction of enzymes involved in the Kennedy (CDP-choline branch) pathway (*Chka, Pcyt1a*) and the Lands’ cycle (*Lpcat3*), which can be traced to Type 1 injured proximal tubular cells [28]. This early activation likely drives enhanced PC biosynthesis and remodeling during the acute injury phase, followed by selective depletion of specific PC species at later time points, explaining the heterogeneous PC profile observed at 24h. In parallel, persistently reduced LPC levels are consistent with efficient early reacylation and limited LPC re-accumulation. Conversely, downregulation of *Pcyt2*, a key enzyme of the Kennedy (CDP-ethanolamine branch) pathway responsible for PE biosynthesis [33], at both 4h and 24h provides a plausible explanation for the sustained reduction in PE levels. Notably, *Pcyt2* is preferentially expressed in healthy proximal tubules and suppressed early after injury, suggesting that PE metabolism is linked to a protective phenotype. Consistent with these results, specific PE (e.g., PE 40:6) and PC species (PC 36:3, PC 38:7, PC 40:6), together with several LPCs, correlated positively with nephroprotective (Klotho, Gpx4) and antioxidant signatures (Cat, Sod1). These associations highlight the potential of selected phospholipid species as biomarkers of tubular repair capacity and therapeutic responsiveness.

Ether phospholipids are integral structural components of cellular membranes, and the presence of alkyl or vinyl ether bonds confers unique biophysical properties that enhance membrane stability and resistance to oxidative stress. Dysregulation of ether phospholipids has been implicated in diverse human diseases, including severe peroxisomal disorders, neurodegenerative and inflammatory diseases, cancer, and metabolic disorders [56,57]. In our study, ether-linked phospholipids (EtherPC and EtherPE) consistently displayed a nephroprotective pattern, positively correlating with established protective markers (*Klotho, Gpx4*) and antioxidant enzymes (*Cat, Sod1*), and inversely with markers of renal dysfunction, tubular injury, and inflammation. Recent studies implicate ether-linked phospholipid biosynthesis in the regulation of lipid peroxidation and ferroptosis, although the functional directionality of these effects remains unresolved [34,35,58,59]. Ferroptosis is an iron-dependent, lipid peroxidation–driven form of regulated cell death that promotes tubular cell loss and exacerbates renal injury in AKI [34,53,59]. Some evidence implicates the peroxisomal ether-lipid synthesis pathway (FAR1–AGPS) in ferroptosis through the generation of ether lipids that serve as substrates for lipid peroxidation [34,60]. In contrast, studies in IRI- and folic acid–induced AKI models report FAR1 downregulation compared with healthy controls [35,53], suggesting a potential protective role in the kidney. Consistently, FAR1 depletion in human proximal tubular cells enhanced ferroptosis markers and oxidative stress, whereas FAR1 overexpression attenuated these responses [35], reinforcing previous evidence that ether lipids protect cells from ferroptotic cell death [61]. However, the role of TMEM189 in plasmalogen synthesis remains debated, with reports describing either no effect or an anti-ferroptotic function [34,60]. GPX4 is a key driver of ferroptosis-mediated renal injury in AKI [62]. Our findings indicate ether-lipid loss was also strongly associated with repression of the *Far1-Agps-Tmem189* axis and *Gpx4*, the central ferroptosis-suppressor enzyme. Accordingly, in a single-cell transcriptomic analyses revealed that *Gpx4* and the *Far1-Agps-Tmem189* axis are predominantly expressed in healthy proximal tubular phenotypes, whereas injured states exhibited coordinated repression of this pathway [28]. Further studies are required to determine whether preservation or therapeutic targeting of ether-lipid metabolism may constitute a promising strategy to limit ferroptosis, oxidative damage, and tubular dysfunction in AKI.

## Conclusions

In conclusion, our holistic approach reveals profound lipid remodeling during IRI and identifies lipid signatures associated with both injury and protection. These findings lay the groundwork for the development of early lipid biomarkers and lipid-targeted therapies aimed at improving outcomes in AKI and kidney transplantation. The strong associations between specific lipid species and either tubular, antioxidant, and ferroptosis-related pathways or kidney protective pathways suggest that lipidomic signatures do not merely reflect tissue injury but capture underlying pathogenic and kidney resilience processes (**Figure 7**). These results are aligned with emerging evidence that lipidomics can outperform traditional biomarkers by detecting metabolic shifts that precede structural damage [15,17]. Future studies should validate these lipid candidates in clinical cohorts and integrate lipidomics with complementary omics technologies to build predictive models capable of forecasting AKI severity, recovery potential, and progression to CKD.

**Figure 7.**
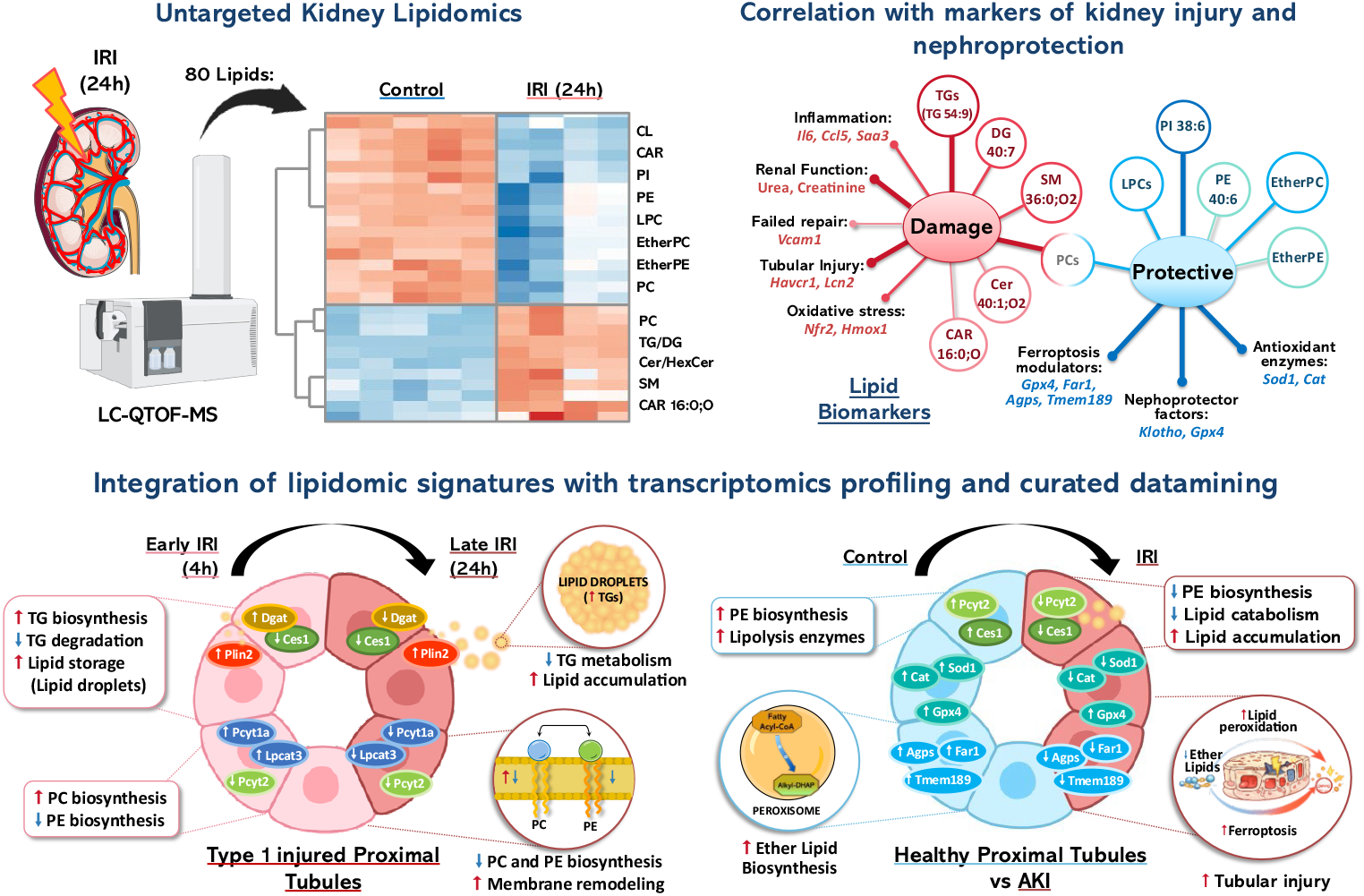
Integrated overview of lipid metabolic reprogramming in kidney ischemia– reperfusion injury (IRI). Untargeted kidney lipidomics by LC-QTOF-MS identified 80 lipid species significantly altered 24 h after IRI, defining distinct lipidomic signatures compared with control kidneys (top left). Correlation analyses revealed damage-associated lipid species, including triglycerides (TGs), diacylglycerols (DGs), sphingolipids, and specific phosphatidylcholines (PCs), which positively correlated with markers of renal dysfunction, tubular injury, inflammation, and oxidative stress, and a nephroprotection-associated lipid module enriched in glycerophospholipid, including PE 40:6, PI 38:6, and several PC/LPC and ether-linked phospholipids, correlating with antioxidant enzymes and nephroprotective factors (top right). Integration with transcriptomic profiling and curated data mining traced triglyceride accumulation, lipid droplet formation, and membrane lipid remodelling to injured proximal tubular cells. In contrast, increased lipid catabolism and the synthesis of phosphatidylethanolamines and ether lipids were correlated with an antioxidant and nephroprotective environment, the loss of which was associated with enhanced tubular injury.

## Abbreviations

Abhd6: Abhydrolase domain containing 6
Acadm: Mitochondrial medium-chain acyl-CoA dehydrogenase
Acox1: Peroxisomal acyl-coenzyme A oxidase 1
Acsl1: Long-chain-fatty-acid--CoA ligase 1
Agps: Peroxisomal alkyldihydroxyacetonephosphate synthase
AKI: Acute kidney injury
CAR: Fatty acyl carnitines
Cat: Catalase
CD36: Platelet glycoprotein 4/ Fatty acid translocase
CDP-DAG: CDP-diacylglycerol
Cept1: Choline/ethanolaminephosphotransferase 1
Ces: Carboxylesterase
CL: Cardiolipin
Cpt1: Carnitine O-palmitoyltransferase 1
Cpt2: Carnitine O-palmitoyltransferase 2
Crls1: Cardiolipin synthase 1
DG (DAG): Diglycerides/Diacylglycerol
Dgat: Diacylglycerol O-acyltransferase
ER: Endoplasmic reticulum
EtherPC: Ether-linked phosphatidylcholine
EtherPE: Ether-linked phosphatidylethanolamine
FA: Fatty acid
FAO: (β-oxidation) Fatty acid β-oxidation
Far1: Fatty acyl-CoA reductase 1
FFAs: Free fatty acids
*Gba2*: Non-lysosomal glucosylceramidase
Gpx4: Glutathione peroxidase 4
HexCer: Hexosylceramide
Hmox1: Heme oxygenase 1
Icam-1: Intercellular adhesion molecule 1
Il6(IL-6): Interleukin 6
IRI: Ischemia–reperfusion injury
IRI-AKI: Ischemia–reperfusion injury–induced acute kidney injury
KIM-1: (Havcr1) Kidney injury molecule 1
KIT: Kidney Interactive Transcriptomics
Lcn2: Neutrophil gelatinase-associated lipocalin
LPC: Monoacyl/diacyl-glycerophosphocholines
Lpcat: Lysophosphatidylcholine acyltransferase
Lpin: Phosphatidate phosphatase
Mgll: Monoglyceride lipase
NGAL: Neutrophil gelatinase-associated lipocalin
Nrf2 (Nfe2l2): Nuclear factor erythroid 2–related factor 2
PA: Phosphatidic acid
PC: Phosphatidylcholine
PE: Phosphatidylethanolamine
PGC-1α: (Ppargc1a) Peroxisome proliferator-activated receptor gamma coactivator 1-alpha
PI: Phosphatidylinositol
Plin2: Perilipin-2
PPARα: Peroxisome proliferator-activated receptor alpha
PS: Phosphatidylserine
Slc27a2: Long-chain fatty acid transport protein 2
SM: Sphingomyelin
Smpd2: Sphingomyelin phosphodiesterase 2
Sod1: Superoxide dismutase 1
Sod2: Superoxide dismutase 2
TG: Triglyceride/triacylglycerol
Tmem189: Transmembrane protein 189
Ugcg: Ceramide glucosyltransferase
Vcam-1: Vascular cell adhesion molecule 1

## ACKNOWLEDGMENTS and Funding

We also gratefully acknowledge Dr. Benjamin Humphreys (Humphreys laboratory) for granting permission to use images from the Kidney Interactive Transcriptomics (KIT) platform. This work was supported by grants from the Instituto de Salud Carlos III (ISCIII) and co-funded by the European Union-NextGenerationEU within the European Union-Plan de Recuperación, Transformación y Resiliencia (MRR) and Fondos Feder (PI23/00394, PI22/00469, PI22/00050, PI21/00251, PI25/00145, ERA-PerMed-JTC2022 (SPAREKID AC22/00027), RICORS program to RICORS2040-renal (RD24/0004/0001; RD24/0004/0021). F.M.S. also acknowledges her Sara Borrell postdoctoral contract (CD23/00049) funded by Instituto de Salud Carlos III (ISCIII) and co-funded by the European Union-NextGenerationEU. This work was also supported by grants from the Comunidad de Madrid Biomedicina to INNOREN-CM “P2022/BMD-7221: Nuevas estrategias diagnósticas y terapéuticas en enfermedad renal crónica” (M.R.O and C.B.) and “P2022/BMD-7223, CIFRA_COR-CM” (A.O.). SPACKDc PMP21/00109, FEDER funds; COST Action PERMEDIK CA21165 supported by COST (European Cooperation in Science and Technology); PREVENTCKD Consortium Project ID 101101220 Programme EU4H DG/Agency HADEA; KitNewCare Project ID 101137054, Call HORIZON-HLTH-2023-CARE-04, Programme HORIZON, DG/Agency HADEA; PICKED Project ID 101168626 HORIZON-MSCA-2023-DN-01-01 MSCA Doctoral Networks 2023.

## REFERENCES

[1] Mehta, R.L. et al. (2015). International Society of Nephrology’s 0by25 initiative for acute kidney injury (zero preventable deaths by 2025): a human rights case for nephrology. The Lancet. 10.1016/S0140-6736(15)60126-X.

[2] Rodríguez, E. et al. (2018). Impact of Recurrent Acute Kidney Injury on Patient Outcomes. Kidney and Blood Pressure Research. 10.1159/000486744.

[3] Rayego-Mateos, S. et al. (2021). Interplay between extracellular matrix components and cellular and molecular mechanisms in kidney fibrosis. Clinical Science. 10.1042/CS20201016.

[4] Ympa, Y.P. et al. (2005). Has mortality from acute renal failure decreased? A systematic review of the literature. The American Journal of Medicine. 10.1016/j.amjmed.2005.01.069.

[5] Siedlecki, A. et al. (2011). Delayed Graft Function in the Kidney Transplant. American Journal of Transplantation. 10.1111/j.1600-6143.2011.03754.x.

[6] Cheval, L. et al. (2024). Lipidomic Profiling of Kidney Cortical Tubule Segments Identifies Lipotypes with Physiological Implications. Function. 10.1093/function/zqae016.

[7] Bhargava, P. and Schnellmann, R.G. (2017). Mitochondrial energetics in the kidney. Nature Reviews Nephrology. 10.1038/nrneph.2017.107.

[8] Bobulescu, I.A. (2010). Renal lipid metabolism and lipotoxicity. Current Opinion in Nephrology and Hypertension. 10.1097/MNH.0b013e32833aa4ac.

[9] Simon, N. and Hertig, A. (2015). Alteration of fatty acid oxidation in tubular epithelial cells: From acute kidney injury to renal fibrogenesis. Frontiers in Medicine. 10.3389/fmed.2015.00052.

[10] Erpicum, P. et al. (2018). What we need to know about lipid-associated injury in case of renal ischemia-reperfusion. American Journal of Physiology - Renal Physiology. 10.1152/ajprenal.00322.2018.

[11] Tammaro, A. et al. (2020). Metabolic Flexibility and Innate Immunity in Renal Ischemia Reperfusion Injury: The Fine Balance Between Adaptive Repair and Tissue Degeneration. Frontiers in immunology. 10.3389/fimmu.2020.01346.

[12] Granata, S. et al. (2022). Oxidative Stress and Ischemia/Reperfusion Injury in Kidney Transplantation: Focus on Ferroptosis, Mitophagy and New Antioxidants. Antioxidants. 10.3390/antiox11040769.

[13] Todorović, Z. et al. (2021). Lipidomics provides new insight into pathogenesis and therapeutic targets of the ischemia—reperfusion injury. International Journal of Molecular Sciences. 10.3390/ijms22062798.

[14] Züllig, T. et al. (2020). Lipidomics from sample preparation to data analysis: a primer. Analytical and Bioanalytical Chemistry. 10.1007/s00216-019-02241-y.

[15] Baek, J. et al. (2022). Lipidomic approaches to dissect dysregulated lipid metabolism in kidney disease. Nature Reviews Nephrology. 10.1038/s41581-021-00488-2.

[16] Cuevas-Delgado, P. et al. (2022). Metabolomics tools for biomarker discovery: applications in chronic kidney disease, in The Detection of Biomarkers, Elsevier, pp. 153–181.

[17] Cuevas-Delgado, P. et al. (2023). Impact of renal tubular Cpt1a overexpression on the kidney metabolome in the folic acid-induced fibrosis mouse model. Frontiers in Molecular Biosciences. 10.3389/fmolb.2023.1161036.

[18] Saiz, M.L. et al. (2024). BET inhibitor nanotherapy halts kidney damage and reduces chronic kidney disease progression after ischemia-reperfusion injury. Biomedicine & Pharmacotherapy. 10.1016/j.biopha.2024.116492.

[19] Cuevas-Delgado, P. et al. (2020). Data-dependent normalization strategies for untargeted metabolomics—a case study. Analytical and Bioanalytical Chemistry. 10.1007/s00216-020-02594-9.

[20] Lange, M. et al. (2021). AdipoAtlas: A reference lipidome for human white adipose tissue. Cell Reports Medicine. 10.1016/j.xcrm.2021.100407.

[21] Gonzalez-Riano, C. et al. (2022). Birth Weight and Early Postnatal Outcomes: Association with the Cord Blood Lipidome. Nutrients. 10.3390/nu14183760.

[22] Gonzalez-Riano, C. et al. (2021). Exploiting the formation of adducts in mobile phases with ammonium fluoride for the enhancement of annotation in liquid chromatography-high resolution mass spectrometry based lipidomics. Journal of Chromatography Open. 10.1016/j.jcoa.2021.100018.

[23] Fernández Requena, B. et al. (2024). LiLA: lipid lung-based ATLAS built through a comprehensive workflow designed for an accurate lipid annotation. Communications Biology. 10.1038/s42003-023-05680-7.

[24] Liebisch, G. et al. (2020). Update on LIPID MAPS classification, nomenclature, and shorthand notation for MS-derived lipid structures. Journal of Lipid Research. 10.1194/jlr.S120001025.

[25] Pang, Z. et al. (2024). MetaboAnalyst 6.0: towards a unified platform for metabolomics data processing, analysis and interpretation. Nucleic Acids Research. 10.1093/nar/gkae253.

[26] Valentijn, F.A. et al. (2021). Ccn2 aggravates the immediate oxidative stress–dna damage response following renal ischemia–reperfusion injury. Antioxidants. 10.3390/antiox10122020.

[27] Szklarczyk, D. et al. (2023). The STRING database in 2023: protein–protein association networks and functional enrichment analyses for any sequenced genome of interest. Nucleic Acids Research. 10.1093/nar/gkac1000.

[28] Li, H. et al. (2022). Comprehensive single-cell transcriptional profiling defines shared and unique epithelial injury responses during kidney fibrosis. Cell Metabolism. 10.1016/j.cmet.2022.09.026.

[29] Nicholson, R.J. et al. (2022). Ceramides and Acute Kidney Injury. Seminars in Nephrology. 10.1016/j.semnephrol.2022.10.007.

[30] Harris, C.A. et al. (2011). DGAT enzymes are required for triacylglycerol synthesis and lipid droplets in adipocytes. Journal of Lipid Research. 10.1194/jlr.M013003.

[31] Braverman, N.E. and Moser, A.B. (2012). Functions of plasmalogen lipids in health and disease. Biochimica et Biophysica Acta (BBA) - Molecular Basis of Disease. 10.1016/j.bbadis.2012.05.008.

[32] Wu, H. et al. (2018). Single-cell transcriptomics of a human kidney allograft biopsy specimen defines a diverse inflammatory response. Journal of the American Society of Nephrology. 10.1681/ASN.2018020125.

[33] van der Veen, J.N. et al. (2017). The critical role of phosphatidylcholine and phosphatidylethanolamine metabolism in health and disease. Biochimica et Biophysica Acta Biomembranes. 10.1016/j.bbamem.2017.04.006.

[34] Cui, W. et al. (2021). Peroxisome-driven ether-linked phospholipids biosynthesis is essential for ferroptosis. Cell Death & Differentiation. 10.1038/s41418-021-00769-0.

[35] Duan, H. et al. (2025). FAR1 as a ferroptosis-related biomarker and potential therapeutic target in acute kidney injury: integrated bioinformatics and experimental validation. Renal Failure. 10.1080/0886022X.2025.2547260.

[36] Zhao, L. et al. (2023). Energy metabolic reprogramming regulates programmed cell death of renal tubular epithelial cells and might serve as a new therapeutic target for acute kidney injury. Frontiers in Cell and Developmental Biology. 10.3389/fcell.2023.1276217.

[37] Rao, S. et al. (2016). Early lipid changes in acute kidney injury using SWATH lipidomics coupled with MALDI tissue imaging. American Journal of Physiology-Renal Physiology. 10.1152/ajprenal.00100.2016.

[38] Scantlebery, A.M.L. et al. (2021). The dysregulation of metabolic pathways and induction of the pentose phosphate pathway in renal ischaemia–reperfusion injury. Journal of Pathology. 10.1002/path.5605.

[39] Rinaldi, A. et al. (2022). Impaired fatty acid metabolism perpetuates lipotoxicity along the transition to chronic kidney injury. JCI Insight. 10.1172/jci.insight.161783.

[40] Li, S.-Y. and Susztak, K. (2018). The Role of Peroxisome Proliferator-Activated Receptor γ Coactivator 1α (PGC-1α) in Kidney Disease. Seminars in Nephrology. 10.1016/j.semnephrol.2018.01.003.

[41] Choi, J. et al. (2024). Carnitine palmitoyltransferase 1 facilitates fatty acid oxidation in a non-cell-autonomous manner. Cell Reports. 10.1016/j.celrep.2024.115006.

[42] Jang, H.-S. et al. (2020). Defective Mitochondrial Fatty Acid Oxidation and Lipotoxicity in Kidney Diseases. Frontiers in Medicine. 10.3389/fmed.2020.00065.

[43] Vasko, R. (2016). Peroxisomes and Kidney Injury. Antioxidants & Redox Signaling. 10.1089/ars.2016.6666.

[44] Johnson, A.C.M. et al. (2005). Triglyceride accumulation in injured renal tubular cells: Alterations in both synthetic and catabolic pathways. Kidney International. 10.1111/j.1523-1755.2005.00325.x.

[45] Zager, R.A. et al. (2011). Acute unilateral ischemic renal injury induces progressive renal inflammation, lipid accumulation, histone modification, and “end-stage” kidney disease. American Journal of Physiology - Renal Physiology. 10.1152/ajprenal.00431.2011.

[46] Zager, R.A. et al. (2005). Renal tubular triglyercide accumulation following endotoxic, toxic, and ischemic injury. Kidney International. 10.1111/j.1523-1755.2005.00061.x.

[47] Kang, H.M. et al. (2015). Defective fatty acid oxidation in renal tubular epithelial cells has a key role in kidney fibrosis development. Nature Medicine. 10.1038/nm.3762.

[48] Asowata, E.O. et al. (2024). Multi-omics and imaging mass cytometry characterization of human kidneys to identify pathways and phenotypes associated with impaired kidney function. Kidney International. 10.1016/j.kint.2024.01.041.

[49] Ruiz-Ortega, M. et al. (2020). Targeting the progression of chronic kidney disease. Nature Reviews Nephrology. 10.1038/s41581-019-0248-y.

[50] Harzandi, A. et al. (2021). Acute kidney injury leading to CKD is associated with a persistence of metabolic dysfunction and hypertriglyceridemia. iScience. 10.1016/j.isci.2021.102046.

[51] Yang, D. et al. (2023). Loss of renal tubular G9a benefits acute kidney injury by lowering focal lipid accumulation via CES1. EMBO reports. 10.15252/embr.202256128.

[52] Nguyen, T.B. et al. (2017). DGAT1-Dependent Lipid Droplet Biogenesis Protects Mitochondrial Function during Starvation-Induced Autophagy. Developmental Cell. 10.1016/j.devcel.2017.06.003.

[53] Martín-Saiz, L. et al. (2022). Ferrostatin-1 modulates dysregulated kidney lipids in acute kidney injury. The Journal of Pathology. 10.1002/path.5882.

[54] Solati, Z. et al. (2018). Oxidized phosphatidylcholines are produced in renal ischemia reperfusion injury. PLOS ONE. 10.1371/journal.pone.0195172.

[55] Marczak, L. et al. (2021). Mass Spectrometry-Based Lipidomics Reveals Differential Changes in the Accumulated Lipid Classes in Chronic Kidney Disease. 10.3390/metabo11050275.

[56] Dean, J.M. and Lodhi, I.J. (2018). Structural and functional roles of ether lipids. Protein & Cell. 10.1007/s13238-017-0423-5.

[57] Bozelli, J.C. et al. (2021). Plasmalogens and Chronic Inflammatory Diseases. Frontiers in Physiology. 10.3389/fphys.2021.730829.

[58] Lee, H. et al. (2021). Ether phospholipids govern ferroptosis. Journal of Genetics and Genomics. 10.1016/j.jgg.2021.05.003.

[59] Kim, J.W. et al. (2023). An integrated view of lipid metabolism in ferroptosis revisited via lipidomic analysis. Experimental & Molecular Medicine. 10.1038/s12276-023-01077-y.

[60] Zou, Y. et al. (2020). Plasticity of ether lipids promotes ferroptosis susceptibility and evasion. Nature. 10.1038/s41586-020-2732-8.

[61] Perez, M.A. et al. (2020). Dietary Lipids Induce Ferroptosis in Caenorhabditiselegans and Human Cancer Cells. Developmental Cell. 10.1016/j.devcel.2020.06.019.

[62] Friedmann Angeli, J.P. et al. (2014). Inactivation of the ferroptosis regulator Gpx4 triggers acute renal failure in mice. Nature Cell Biology. 10.1038/ncb3064.

